# Discovery of a Potent and Dual-Selective Bisubstrate Inhibitor for Protein Arginine Methyltransferase 4/5

**DOI:** 10.1101/2020.09.05.283978

**Authors:** Ayad A. Al-Hamashi, Dongxing Chen, Youchao Deng, Guangping Dong, Rong Huang

**Author notes:** Address correspondence to this author at the Department of Medicinal Chemistry and Molecular Pharmacology, Purdue University, 720 Clinic Drive, West Lafayette, Indiana, 47907, USA.

## Abstract

Protein arginine methyltransferases (PRMTs) have been implicated in the progression of many diseases. Understanding substrate recognition and specificity of individual PRMT would facilitate the discovery of selective inhibitors towards future drug discovery. Herein, we reported the design and synthesis of bisubstrate analogues for PRMTs that incorporate a S-adenosylmethionine (SAM) analogue moiety and a tripeptide through an alkyl substituted guanidino group. Compound **AH237** is a potent and selective inhibitor for PRMT4 and PRMT5 with a half-maximal inhibition concentration (IC_50_) of 2.8 nM and <1.5 nM, respectively. Computational studies provided a plausible explanation for the high potency and selectivity of **AH237** for PRMT4/5 over other 40 methyltransferases. This proof-of-principle study outlines an applicable strategy to develop potent and selective bisubstrate inhibitors for PRMTs, providing valuable probes for future structural studies.

## 1. Introduction

Arginine methylation is a prominent post-translational modification that regulates various physiological processes including cell differentiation, RNA splicing, and DNA damage repair.^1^ It is catalyzed by protein arginine methyltransferases (PRMTs) that transfer the methyl group from the cofactor S-adenosylmethionine (SAM) to the guanidino group of the arginine residue in an “SN2-like” fashion. There are nine members in the PRMT family, which are categorized into three types based on the product types.^1,2^ Type I PRMTs (PRMT1, 2, 3, 4, 6, and 8) generate asymmetric dimethylated arginine. Type II PRMTs (PRMT5 and 9) produce symmetric dimethylated arginine. Type III contains only PRMT7 that yields monomethylated arginine. Besides a typical Rossmann-fold domain for interaction with the cofactor SAM, most of PRMTs share the glycine and arginine (GAR) substrate motif.^3^ The only exception is PRMT9 which displays the preference for FKRKY motif.^4^ Although PRMT4 predominantly recognize arginine residues in proline-rich context,^5^ it is able to methylate arginine residues next to proline, glycine, and methionine rich motifs in vitro. Therefore, it remains ambiguous about the physiological substrate preference for each PRMT.

Abnormal expression or activity of PRMTs has been associated with a variety of diseases, including cancers, cardiovascular diseases, inflammatory diseases, and diabetes.^6–10^ Thus, PRMTs have attracted emerging attentions to develop specific and potent inhibitors as potential therapeutic agents. Although many potent small molecule inhibitors have been reported for PRMTs to date, selectivity remains a challenge for individual PRMT isoform because of a conserved SAM binding site and similar substrate recognition motif.^9,11–14^ Bisubstrate analogue has demonstrated its potential to offer potent and specific inhibitors for several PRMTs including PRMT1, PRMT4, and PRMT6, as well as facilitate the formation of co-crystal structures to offer valuable insights into the structural basis for PRMT specificity.^15–17^ For instance, the bisubstrate inhibitor **GMS** that links a guanidine moiety to a sinofungin (SNF) analogue is 17-fold more potent than SNF for PRMT6 (Figure 1).^15^ Similarly, bisubstrate analogue **MH4** that tethers an adenosine moiety with the PRMT4 substrate peptide PAAPRPPFSTM displays potent inhibition of PRMT4 with IC_50_ value of 90 nM and 250-fold selectivity over PRMT1.^17^ **JNJ-64619178** is a highly selective PRMT5 inhibitor in phase I clinical trial for advanced solid tumors, non-Hodgkin lymphoma, and myolodysplastic syndromes.^18^ In this study, we designed a general platform to develop potent and selective inhibitors for PRMTs. We adopted both rational and structure-based drug design strategies to produce a series of new bisubstrate analogues that covalently connect a SAM analogue with a single amino acid (Lys or Arg) or a tripeptide (RGR or RGK) through a guanidine group (Figure 2), which is the methyl acceptor of arginine. Among them, **AH237** showed a superior selectivity for PRMT4 and PRMT5 with IC_50_ values in a low nanomolar range.

**Figure 1.**
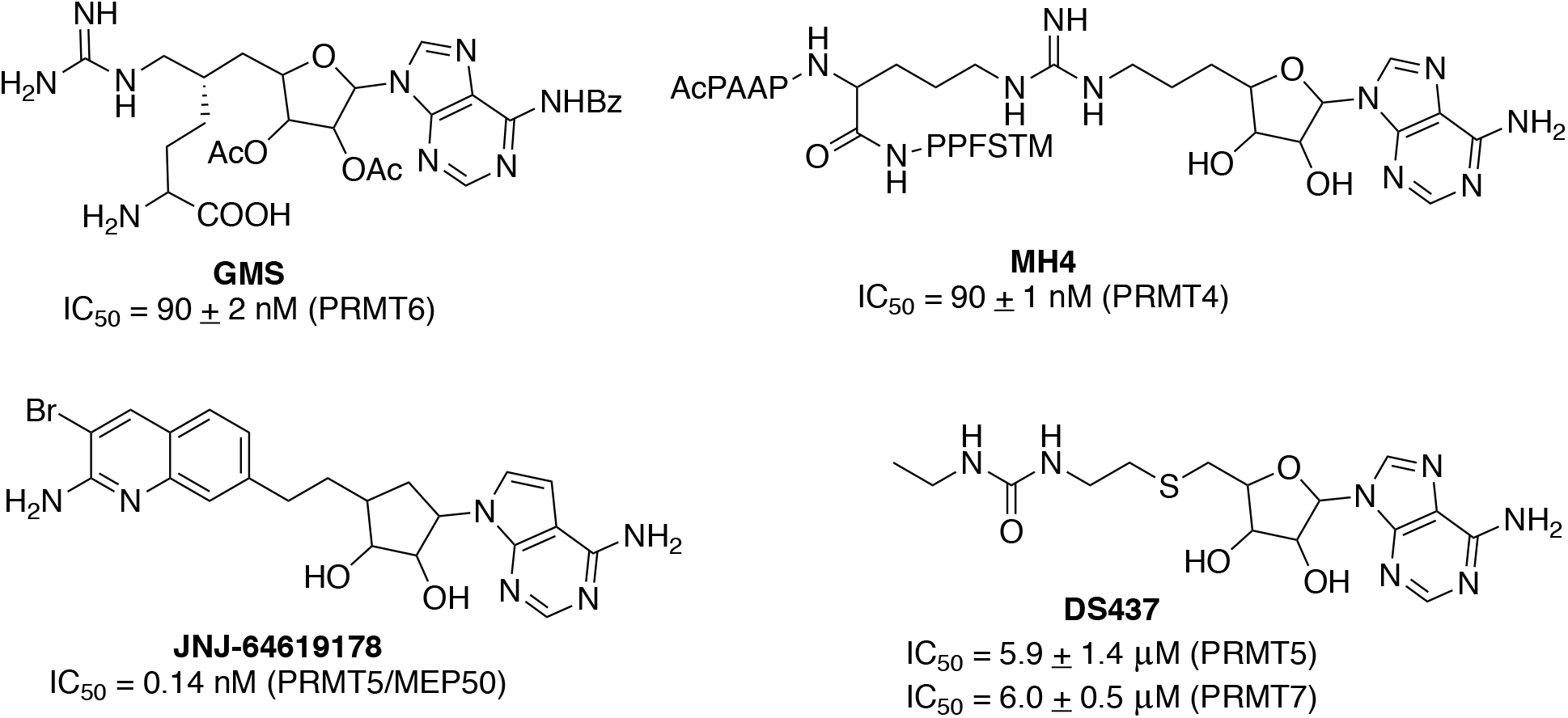
Reported bisubstrate inhibitors for PRMTs.

**Figure 2.**
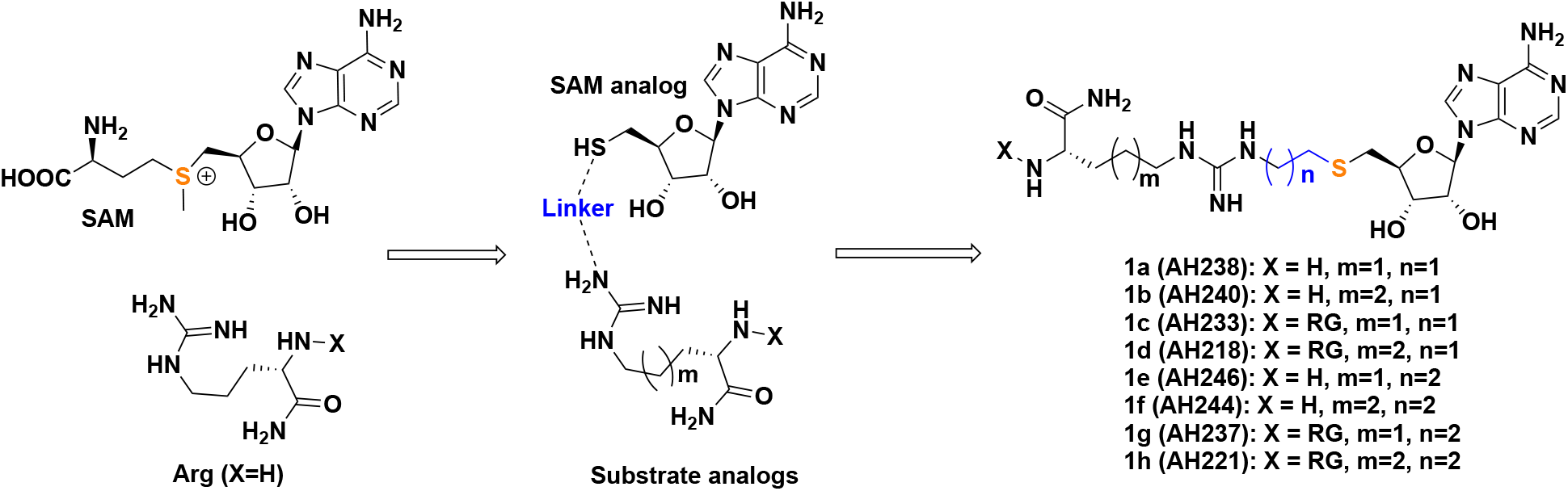
The designed PRMT inhibitors.

## 2. Results and Discussion

### 2.1. Design

In the active site of PRMTs, two adjacent binding pockets are occupied by both S-(5’-Adenosyl)-L-homocysteine (SAH) and the Arg residue of the substrate peptide. The distance between the SAH sulfur atom and the α-nitrogen atom of the guanidinum group ranges from 3.5 Å to 5.5 Å.^15–17^ Therefore, we hypothesized that covalently linking a SAM analogue moiety with a guanidinum group through a 2-C or 3-C atom linker would mimic the transition state to offer potent inhibitors as general probes for PRMTs (Figure 2). A 2-C or 3-C atom linker length has demonstrated feasibility for various protein methyltransferases in previous studies.^15–17,19–22^ For the SAM cofactor analogue, we chose a thioadenosine as inspired by a dual PRMT5/7 inhibitor **DS437**.^23^ For the substrate portion, we focused on a general peptide motif RGR since it can be essentially recognized by all PRMTs except PRMT9. To achieve the specificity for PRMTs, the methylation acceptor guanidium group was retained. We also investigated an RGK peptide, as well as a single amino acid (Arg or Lys), to explore the effect of substrate peptide moiety on the inhibition (Table 1 and Figure 2).

**Table 1.** The inhibitory IC_50_ of the synthesized compounds.

| ID | R1 | n | IC<sub>50</sub> (μM) |  |  |  |  |
| --- | --- | --- | --- | --- | --- | --- | --- |
|  |  |  | PRMT1<sup>a</sup> | TbPRMT7<sup>a</sup> | G9a<sup>b</sup> | NTMT1<sup>b</sup> | SETD7<sup>b</sup> |
| **1a (AH238)** | Orn | 1 | 16±8 | 2.8±0.2 | >100 | >100 | >100 |
| **1b (AH240)** | Lys | 1 | 8.3±5.4 | 3.3±2.2 | >100 | >100 | >100 |
| **1c (AH233)** | Arg-Gly-Orn | 1 | 1.7±0.1 | 0.59±0.05 | >100 | >100 | >100 |
| **1d (AH218)** | Arg-Gly-Lys | 1 | 8.3±0.1 | 0.91±0.07 | ~10 | >100 | ~100 |
| **1e (AH246)** | Orn | 2 | 6.5±2.8 | 7.6±1.0 | >100 | >100 | >100 |
| **1f (AH244)** | Lys | 2 | 3.0±1.1 | 20±6 | >100 | >100 | >100 |
| **1g (AH237)** | Arg-Gly-Orn | 2 | 15±0.2 | 0.29±0.03 | >100 | >100 | >100 |
| **1h (AH221)** | Arg-Gly-Lys | 2 | 5.6±0.1 | 3.4±0.3 | >100 | >100 | >100 |

Note: <sup>a</sup> IC<sub>50</sub> values were determined in triplicate (n= 3) and presented as mean±standard deviation (SD). <sup>b</sup> IC<sub>50</sub> values were estimated from three-dose screening in duplicate (n=2).

### 2.2. Synthesis

Commercially available adenosine was first subjected to a diol protection and followed by a subsequent Mistunobu reaction to produce thioester **2** (Scheme 1).^19,24^ Then **2** was hydrolyzed by sodium methoxide and followed by thiol alkylation with phthalimide alkyl bromides **3a-b** to provide phthalimides **4a-b**.^25^ Deprotection of **4a-b** by hydrazine afforded the key amine intermediates **5a-b**, which were then reacted with various thioureas **6a-d** to provide **7a-h** in a convergent manner.^26,27^ Removal of Fmoc protection group followed by acidic deprotection or cleavage offered final compounds **1a-h**.^28^ For **1a-b** and **1e-f**, Fmoc-Orn(Mtt) and Fmoc-Lys(Boc) were first amidated using ammonium chloride to produce **9a-b**, followed by deprotection with TFA to yield free amines to react with Fmoc-isothiocyanate at 0°C to yield **6a-b**.^27^ To prepare peptide conjugates **1c-d** and **1g-h**, short peptides Fmoc-Arg-Gly-Orn/Lys (**9a-b**) were synthesized on solid phase and followed by removal of Mtt protection group, which were then reacted with Fmoc-isothiocyanate to yield **6c-d**.^27^

**Scheme 1.**
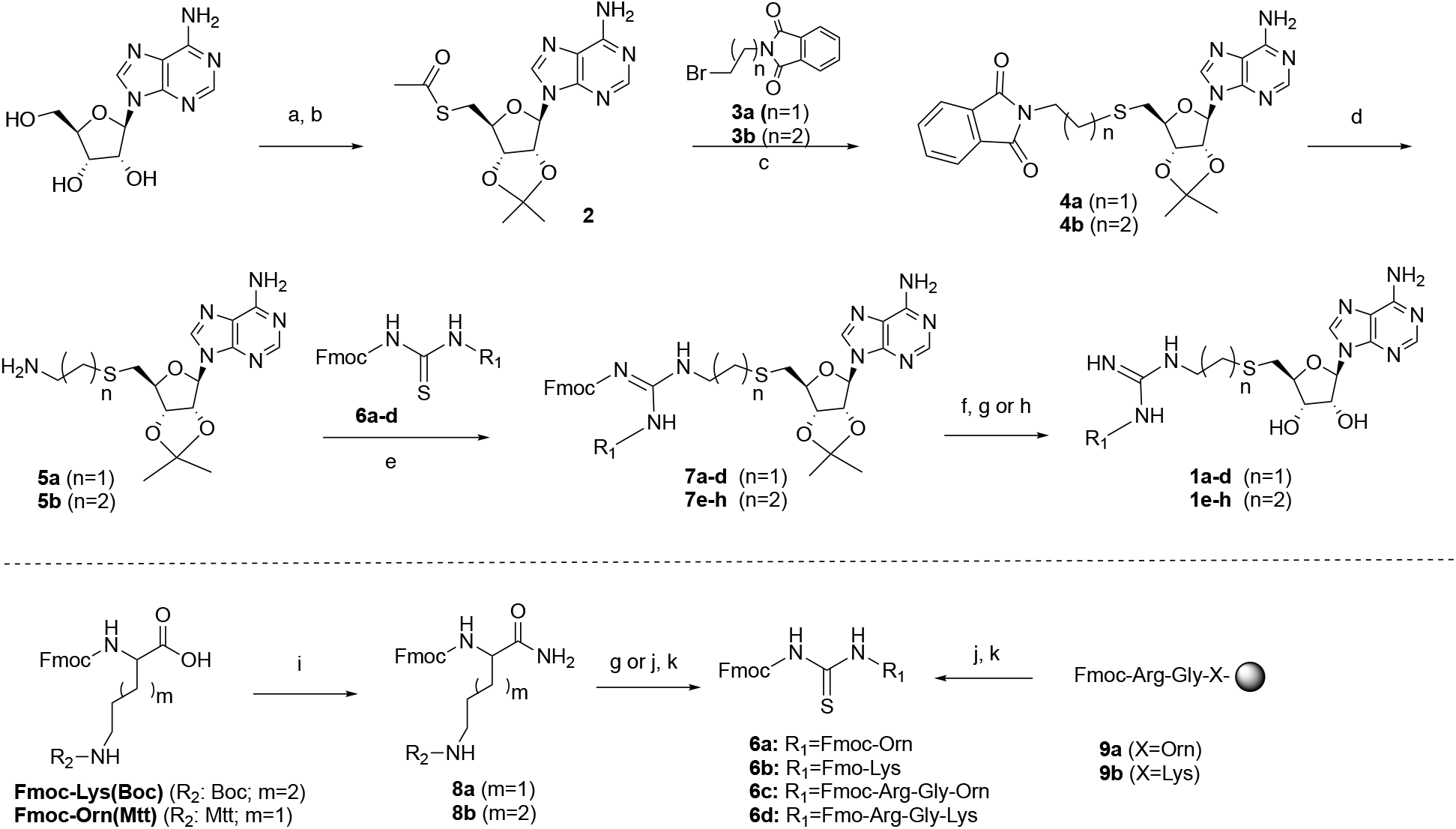
Synthesis of compounds **1a-h**. Reagents and conditions: (a) CH(OEt)_3_, p-TsOH, acetone, 85%; (b) Thioacetic acid, PPh_3_, DIAD, THF, 94%; (c) NaOCH_3_, MeOH, 59-70%; (d) Hydrazine, MeOH, 76-92%; (e) EDC, DIPEA, CH_2_Cl_2_; (f) Piperidine, HOBt, DMF; (g) 20% TFA in CH_2_Cl_2_, 0 °C; (h) TFA:DODT:TIPS:H_2_O (94:2.5:1:2.5 v/v), rt, 5 h; (i) NH_4_Cl, HBTU, NMM, CH_3_CN, 70-90%; (j) 1 % TFA in CH_2_Cl_2_; (k) Fmoc-isothiocyanate, CH_2_Cl_2_, 0 °C.

### 2.3. Biochemical Characterization

All synthesized bisubstrate analogues were first evaluated in a SAH hydrolase (SAHH)-coupled fluorescence assay under the condition of the *K*_m_ values of both SAM and the respective peptide substrate for two representative PRMTs (PRMT1 and *Tb*PRMT7).^21,29^ As shown in Table 1 and Figure 3A-B, all bisubstrate analogues exhibited inhibition against PRMT1 and *Tb*PRMT7 with IC_50_ values ranging from 1.7 to 14.6 μM and 0.29 to 19.5 μM, respectively. The majority of bisubstrate analogues displayed improved or comparable potency to *Tb*PRMT7 than PRMT1 except **AH244**, which showed 6-fold increased potency to PRMT1. According to results from this series, PRMT1 demonstrated its preference to a 3-C atom linker while *Tb*PRMT7 showed its preference for a 2-C atom linker. For instance, **AH244** and **AH246** that contained a 3-C atom linker were around 3-fold and 14-fold more potent for PRMT1 than **AH240** and **AH238** that contained a 2-C atom linker, respectively. However, **AH233** and **AH237** were exceptions because **AH237** containing a 3-C atom linker was the most potent inhibitor (IC_50_ = 0.29 μM) for *Tb*PRMT7 and **AH233** containing a 2-C atom linker was the most potent inhibitor (IC_50_ = 0.59 μM) for PRMT1 in this series. Furthermore, **AH237** displayed over 50-fold selectivity for *Tb*PRMT7 over PRMT1. In terms of the effect of peptide length on inhibition, bisubstrate analogues containing a tripeptide substrate showed about 4-fold improved potency for *Tb*PRMT7 in comparison with their respective bisubstrate analogues containing a single amino acid, while marginal effects were observed for PRMT1 except for **AH233**. Meanwhile, the linker length has minimal effect on the bisubstrate analogues that contain the tripeptide moiety like Arg-Gly-Arg for *Tb*PRMT7 as shown in **AH233** and **AH237**, but about 8-fold difference for PRMT1.

**Figure 3.**
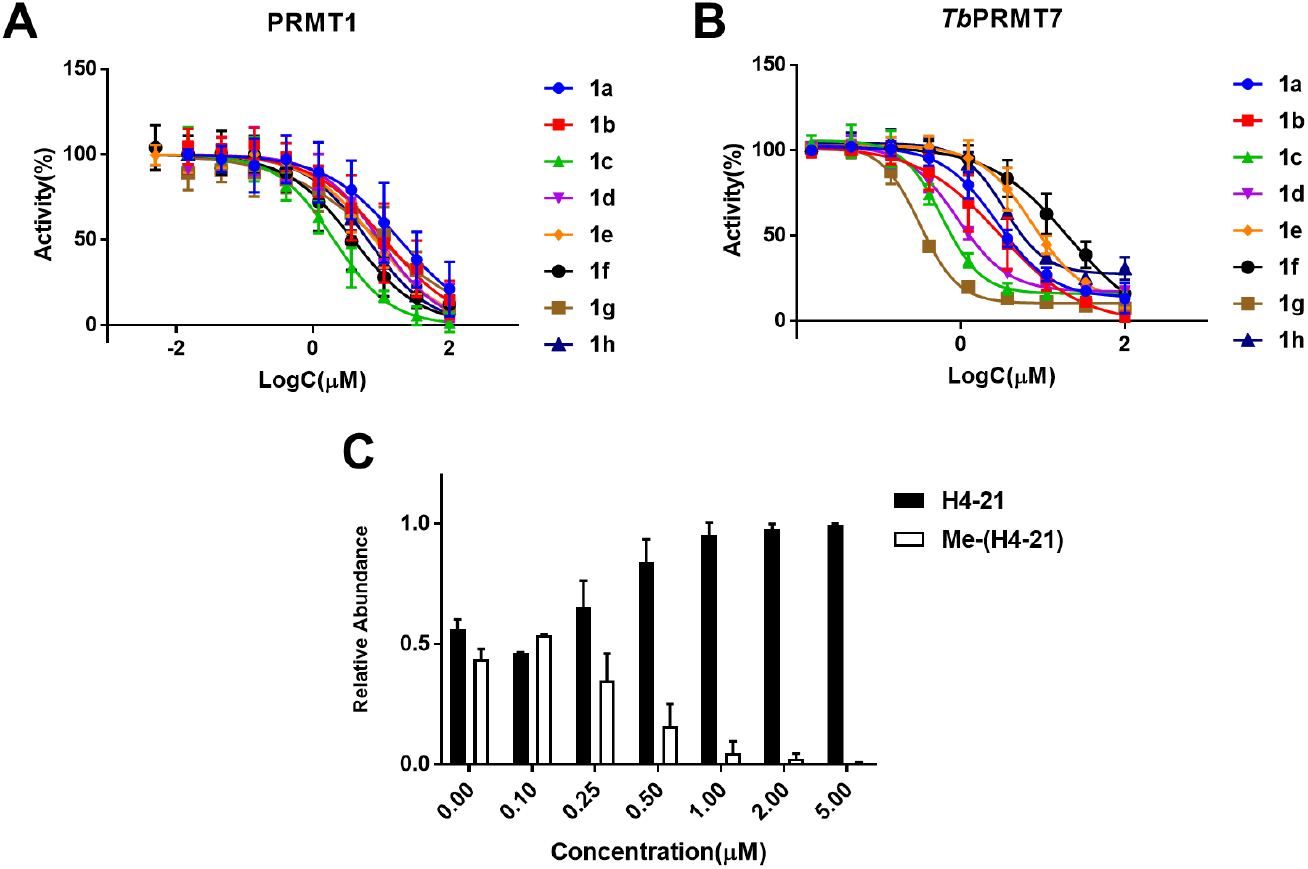
(A) IC_50_ curves of bisubstrate compounds for PRMT1. (B) IC_50_ curves of bisubstrate compounds for *Tb*PRMT7. (C) The methylation assay of **AH237** for *Tb*PRMT7 (n=2).

Our hypothesis is that designed PRMT inhibitors would be selective for PRMTs against other protein methyltransferases such as protein lysine methyltransferases (PKMTs) and N-terminal methyltransferases (NTMTs), because all designed compounds contain a guandinium function group that is a unique methylation acceptor for PRMTs. To test this hypothesis, we chose two representative PKMTs (G9a and SETD7) and NTMT1 to examine their activities in the SAHH-coupled fluorescence assay.^29^ Not surprisingly, all eight bisubstrate analogues did not display any significant inhibition against G9a, NTMT1, and SETD7 up to 100 μM, except that **AH218** inhibited 50% of G9a activity at 10 μM. **AH237** that incorporated a tripeptide of Arg-Gly-Arg and a 3-C atom linker was the most potent and selective inhibitor for *Tb*PRMT7 in our series (IC_50_ = 0.29 ± 0.03 μM). When we carried out this study, no potent and selective PRMT7 inhibitor was available except a dual PRMT5 and 7 inhibitor **DS437** (IC_50_ = 6 μM).^23^ Therefore, we focused on **AH237** in subsequent inhibition mechanism and the comprehensive selectivity study.

### 2.4. MALDI-MS Methylation Inhibition Assay

The MALDI-MS methylation assay was performed to validate the inhibitory activity of **AH237** on *Tb*PRMT7.^30,31^ The results indicated that at 0.5 μM of **AH237** the methylation level of H4-21 was reduced by more than 50%, while the methylated product was abolished with 5 μM compound (Figure 3C).

### 2.5. Inhibition Mechanism

To examine the inhibition mechanism of **AH237**, a kinetic analysis was performed using the SAHH-coupled fluorescence-based assay with *Tb*PRMT7. **AH237** showed an unambiguous pattern of competitive inhibition for both the peptide substrate and SAM, as demonstrated by the linear ascending of the IC_50_ values depending on either the peptide substrate or SAM concentration (Figure 4). The result indicated that compound **AH237** occupied both cofactor and peptide substrate binding sites of *Tb*PRMT7 as a bisbustrate inhibitor, supporting its feature as the bisubstrate analogue.

**Figure 4.**
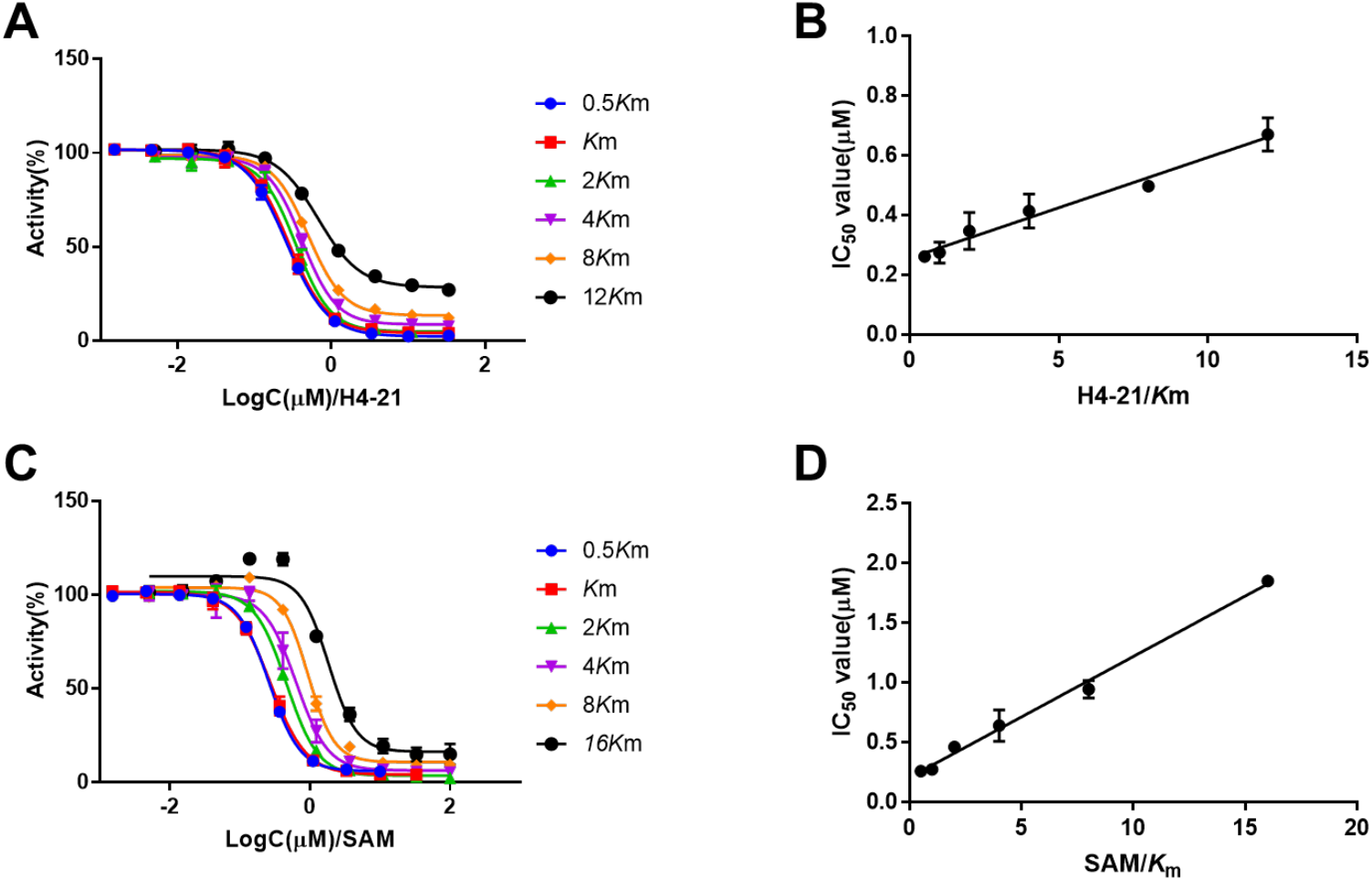
Inhibition mechanism studies of **AH237** for *Tb*PRMT7. (A) IC_50_ curves of **AH237** at varying concentrations of H4-21 peptide with fixed concentration of SAM. (B) Linear regression plot IC_50_ values with corresponding concentrations of H4-21. (C) IC_50_ curves of **AH237** at varying concentrations of SAM with fixed concentration of H4-21. (D) Linear regression plot IC_50_ values with corresponding concentrations of SAM.

### 2.6. Selectivity Studies

To further understand the selectivity profile of **AH237**, its inhibitory activity was examined for a panel of 41 MTases such as PRMTs (PRMT1, 3-8), PKMTs (ASH1L, EZHs, G9a, GLP, METTL21A, MLL complexes, NSDs, PRDM9, SETs, SMYDs, DOT1L, and SUV39Hs), NRMT1/2, and DNA methyltransferases (DNMTs) at a single dose (10 μM) of **AH237** (Reaction Biology Inc.). The results indicated that **AH237** selectively reduced the activity for most PRMTs at 10 μM except PRMT3. Importantly, **AH237** did not show any inhibition for all PKMTs, NTMTs, and DNMTs except for SMYD2 and 3 (Figure 5, Table S1). To our surprise, **AH237** completely abolished the activities of PRMT4 and 5 at the concentration of 10 μM. Subsequent dose-response analysis indicated that **AH237** is highly selective for PRMT4 and PRMT5 with IC_50_s of 2.8±0.17 nM and <1.5 nM, respectively. Its potency was also confirmed for PRMT1 (IC_50_ = 5.9±2.3 μM) and PRMT7 (IC_50_=831±93 nM), which are comparable to the values in Table 1 (Figure 6).

**Figure 5.**
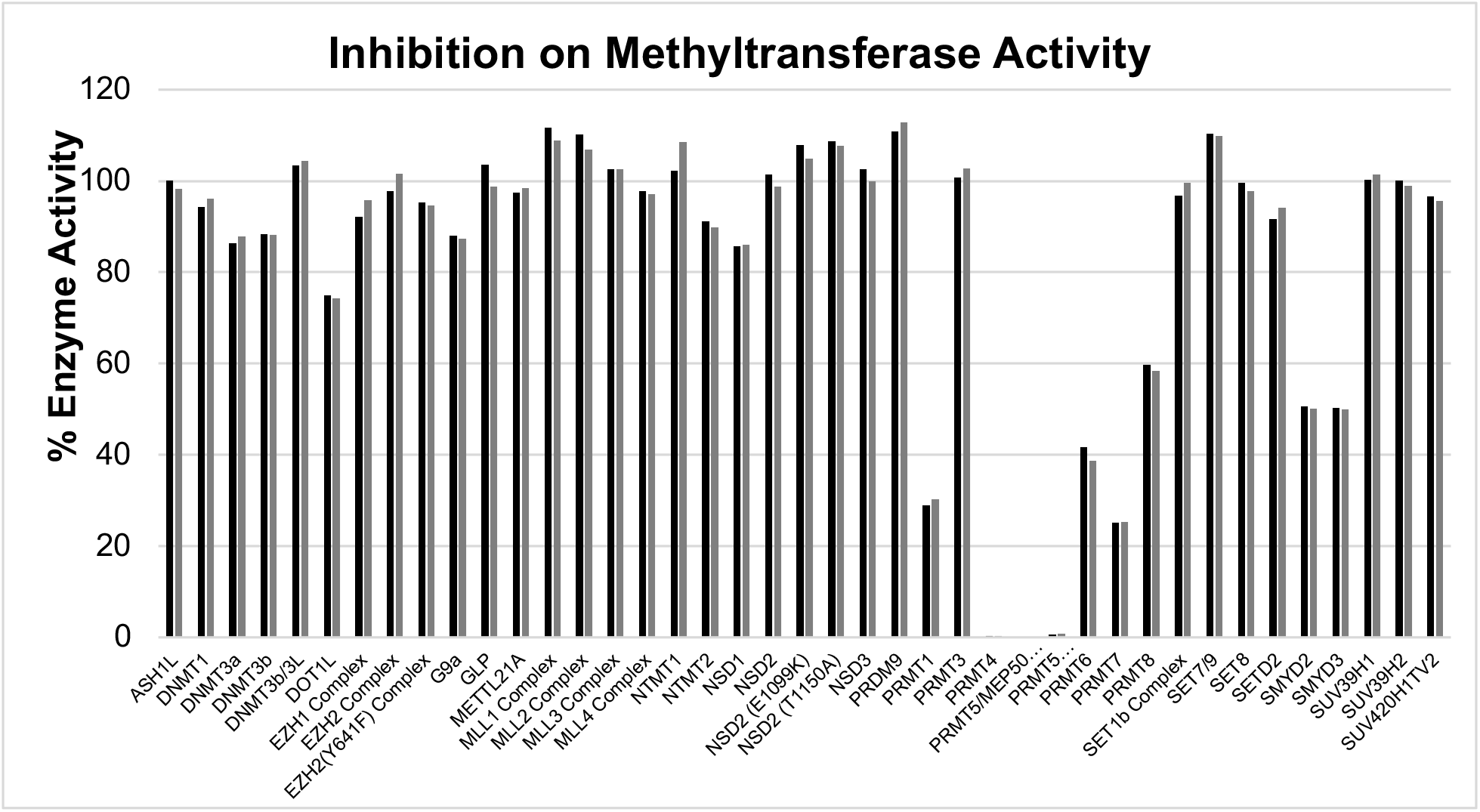
HotSpot methyltransferases inhibition profile of **AH237** at 10 μM on various methyltransferase enzymes in duplicates (n=2). The results indicated that PRMT enzymes, including PRMT4 and PRMT5, were selectively inhibited by **AH237** except PRMT3.

**Figure 6.**
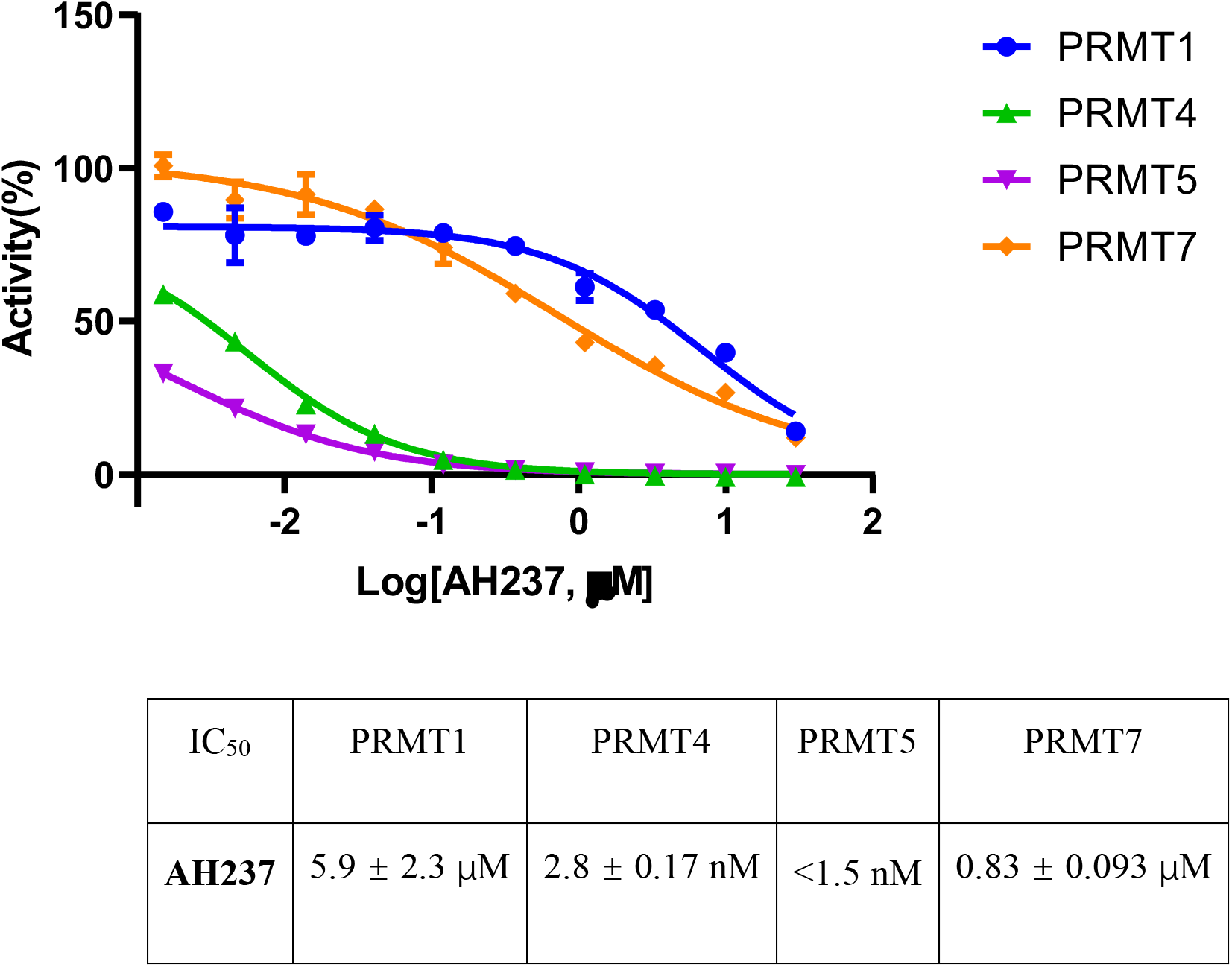
IC_50_ determination of **AH237** against human PRMT1/4/5/7 (n=2).

### 2.7. Docking Studies

In an attempt to rationalize the observed high potency and selectivity of **AH237** for PRMT4/5, we performed computational studies for PRMT1, 4, 5, and 7. Interactions of thioadenosine moiety of **AH237** with those four PRMTs offered plausible explanation for its different inhibitory activities towards PRMT1/4/5/7. As shown in Figure 7, the thioadenosine moiety of **AH237** overlaid very well with SAH or SAH mimic moiety in PRMT4 and 5.^17,32^ For PRMT7, most interactions were retained except a slight shift, which possibly resulted in the loss of interaction with Glu125.^33^ Whereas, the interaction patterns of **AH237** were quite different for PRMT1 compared to the interaction of SAH with PRMT1, yielding loss of the interactions with both Cys101 and Glu100 of PRMT1.^34^ Together, these results provided insight into the preferences of **AH237** binding to PRMT4/5 over PRMT1/7. In addition to the adenosine moiety, the tripeptide portion of **AH237** demonstrated very similar binding orientation as the peptide substrate of PRMT4/5. Furthermore, the linker guanidine group formed more interactions with PRMT4/5 than PRMT7 (Figure 8).

**Figure 7.**
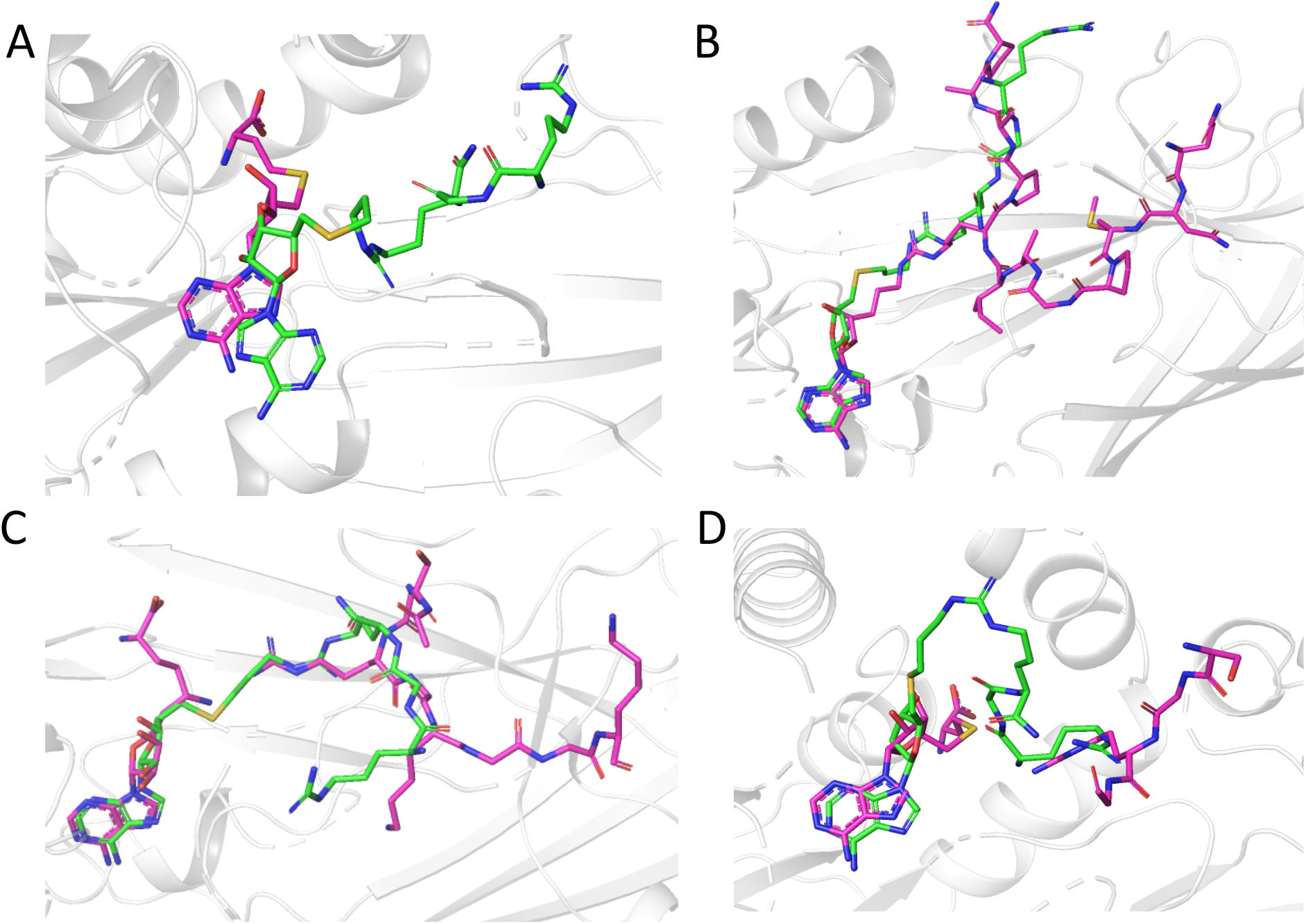
Predicted binding modes of **AH237** in the PRMTs active site. **AH237** was shown in green stick model, while peptides, SAM or SAH were displayed with magenta stick model. Residues in PRMTs were hidden for a better view. (A) PRMT1 (PDB, 1OR8). (B) PRMT4 (PDB, 5LGP). (C) PRMT5 (PDB, 4GQB). (D) PRMT7 (PDB, 4M38).

**Figure 8.**
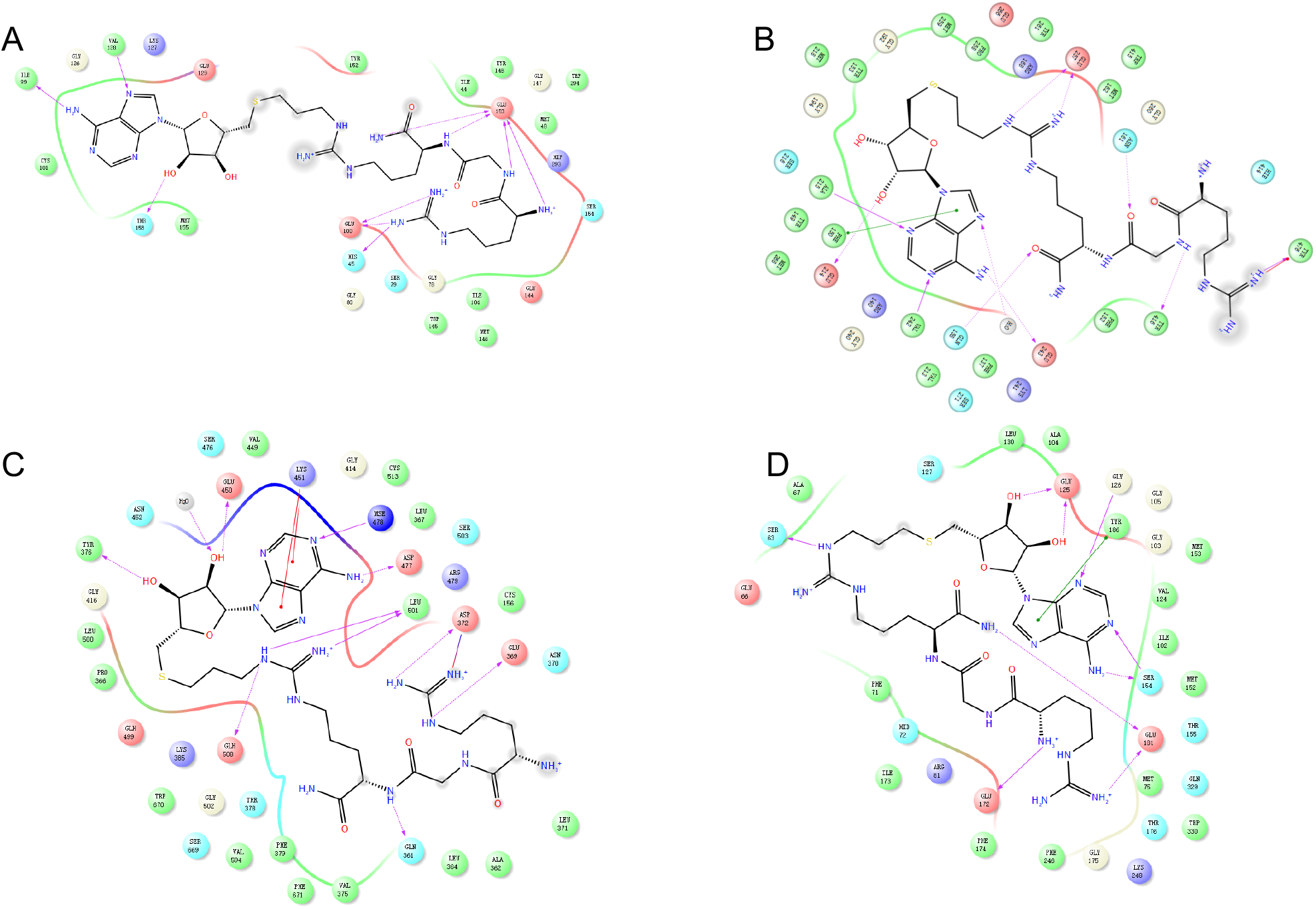
The estimated 2D interactions of **AH237** in the pocket of PRMTs. The peptides, SAM, or SAH in PRMTs were hidden for a better view. **AH237** was shown in black line model, while residues around the protein pocket were presented as colored circles. (A) PRMT1 (PDB, 1OR8). (B) PRMT4 (PDB, 5LGP). (C) PRMT5 (PDB, 4GQB). (D) PRMT7 (PDB, 4M38).

## 3. Conclusion

In summary, we have designed and synthesized a series of new bisubstrate inhibitors for PRMTs that covalently link either a single amino or peptide with a thioadenosine through a guanidino group. All of the synthesized inhibitors showed selectivity for PRMTs over two representative PKMTs (G9a and SETD7) and NTMT1. In general, PRMT1 showed less sensitivity to bisubstrate analogues with variable linkers and substrate sequence length because there is less than a 10-fold difference among eight bisubstrate analogues. This tolerance supports a broad substrate spectrum of PRMT1 as it mediates over 85% of the reported arginine methylation events.^10^ On the contrary, PRMT7 exhibited higher stringency towards substrate moiety as bisubstrate analogues containing a tripeptide portion were more potent for *Tb*PRMT7 than their respective pair with a single amino acid.^35^ Among them, **AH237** showed high potency for PRMT4 and 5 with an IC_50_ value of 2.8 nM and less than 1.5 nM, respectively. Moreover, it displayed almost 1,000-fold selectivity over PRMT1 and 7, and over 10,000-fold selectivity for the other MTases. This profound selectivity corroborates the benefits of bisubstrate analogues that are able to differentiate their potency even among PRMTs family that share a similar substrate recognition preference. In summary, our study offers a glimpse of a delicate difference of transition state for PRMTs, which shed lights on the specificity of PRMT4/5 over the other PRMTs. Additionally, our study outlines a feasible and applicable synthetic strategy to develop bisubstrate inhibitors for PRMTs.

## 4. Experimental

### 4.1. Material and Instruments

All chemicals and solvents were purchased from commercial suppliers and used without further purification unless stated otherwise. ^1^H and ^13^C-NMR spectra were carried out on Brucker Avance 500 MHz NMR spectrometer in deuterated solvents. High-resolution Matrix-assisted laser desorption/ionization (MALDI) spectra were performed on 4800 MALDI TOF/TOF mass spectrometry (Sciex) at the Mass Spectrometry and Purdue Proteomics Facility (PPF), Purdue University. Peptides were synthesized by CEM Liberty Blue peptide synthesizer. Crude products were purified by chromatography using silica gel, standard grade from DAVSIL^®^ (code number 1000179164, 35-70 micron SC). Flash chromatography was performed on Teledyne ISCO CombiFlash Companion chromatography system on RediSep prepacked silica cartridges. Thin-layer chromatography (TLC) plates (20 cm x 20 cm) were purchased from Merck KGaA.

### 4.2. Synthesis

#### 4.2.1. Synthesis *S*-(((3a*S*,4*S*,6*R*,6a*R*)-6-(6-Amino-9*H*-purin-9-yl)-2,2-dimethyltetrahydrofuro[3,4-*d*] [1,3]dioxol-4-yl)methyl) ethanethioate (2)

To a solution of the adenosine (0.8 g, 3 mmol, 1 equiv) in acetone (100 mL) was added triethylorthoformate (2.9 g, 19.5 mmol, 6.5 equiv), and p-toluenesulfonic acid (2.55 g, 15 mmol, 5 equiv). The reaction mixture was stirred at room temperature overnight, then was quenched with sat. NaHCO_3_ (50 mL) and extracted with ethylacetate (3 x 40 mL). The combined organic extract was washed with brine, dried over anhydrous sodium sulfate, and the solvent was evaporated under *vacuo* to get (2.5 mmol) protected adenosine. Then the product was dissolved in tetrahydrofuran (40mL) and added triphenyl phosphine (2 g, 7.5 mmol, 3 equiv), DIAD (1 g, 5 mmol, 2 equiv), and thioacetic acid (395 mg, 5 mmol, 2 equiv). The reaction mixture was stirred at room temperature for 2 hr. It was concentrated in *vacuo* and purified by flash chromatography on silica gel in 60-90% acetone/hexanes to get **2** as a fluffy white solid (875 mg, 80%, in two steps). ^1^H NMR (500 MHz, CDCl_3_) δ 8.36 (s, 1H), 8.17 (s, 1H), 6.16 (d, *J* = 2.4 Hz, 1H), 5.31 – 5.26 (m, 1H), 4.94 (dd, *J* = 6.3, 3.2 Hz, 1H), 4.58 (dt, *J* = 5.5, 3.7 Hz, 1H), 4.35 – 4.25 (m, 2H), 1.99 (s, 3H), 1.66 – 1.63 (s, 3H), 1.42 – 1.39 (s, 3H). ^13^C NMR (126 MHz, CDCl_3_) δ 194.58, 155.72, 152.77, 149.09, 140.02, 120.24, 114.52, 90.90, 86.13, 84.20, 83.68, 31.26, 30.57, 27.07, 25.35.

#### 4.2.2. General procedure for synthesis of phthalidomides 4a-b

To a solution of **2** (730 mg, 2 mmol, 1 equiv) and bromoalkyl phthalidomides **3a-b** (4 mmol, 2 equiv) in methanol (50 mL) at -20 °C was gradually added sodium methoxide (30% w/w in methanol) (0.55 ml, 3 mmol, 1.5 equiv). The reaction mixture was stirred at -20 °C for 3 hr and then at room temperature overnight. H_2_O was added (20 ml) and extracted with ethylacetate (3 x 50 mL). The combined organic extract was washed with brine, dried over anhydrous sodium sulfate, and the solvent was evaporated under *vacuo* and purified by flash chromatography on silica gel in 60-90% acetone/hexanes.

##### 4.2.2.1. 2-(2-((((3a*S*,4*S*,6*R*,6a*R*)-6-(6-Amino-9*H*-purin-9-yl)-2,2-dimethyltetrahydrofuro[3,4-*d*] [1,3]dioxol-4-yl)methyl)thio)ethyl)isoindoline-1,3-dione (4a)

^1^H NMR (500 MHz, CDCl_3_) δ 8.36 (s, 1H), 7.97 (s, 1H), 7.82 (dd, *J* = 5.4, 3.0 Hz, 2H), 7.69 (dd, *J* = 5.5, 3.0 Hz, 2H), 6.31 (s, 2H), 6.08 (d, *J* = 2.3 Hz, 1H), 5.45 (dd, *J* = 6.4, 2.3 Hz, 1H), 5.05 (dd, *J* = 6.4, 3.3 Hz, 1H), 4.40 (td, *J* = 6.7, 3.4 Hz, 1H), 3.80 (t, *J* = 7.3 Hz, 2H), 2.96 – 2.86 (m, 2H), 2.81 (t, *J* = 7.0 Hz, 2H), 1.59 (s, 3H), 1.37 (s, 3H).

##### 4.2.2.2. 2-(3-((((3a*S*,4*S*,6*R*,6a*R*)-6-(6-Amino-9*H*-purin-9-yl)-2,2-dimethyltetrahydrofuro[3,4-*d*|[1,3]dioxol-4-yl)methyl)thio)propyl)isoindoline-1,3-dione (4b)

^1^H NMR (500 MHz, CDCl_3_) δ 8.33 (s, 1H), 7.96 (s, 1H), 7.85 – 7.78 (m, 2H), 7.72 – 7.66 (m, 2H), 6.25 (s, 2H), 6.07 (d, *J* = 2.2 Hz, 1H), 5.46 (dd, *J* = 6.4, 2.3 Hz, 1H), 5.03 (dd, *J* = 6.4, 3.2 Hz, 1H), 4.42 – 4.34 (m, 1H), 3.72 (t, *J* = 7.4 Hz, 3H), 2.90 – 2.72 (m, 2H), 2.55 (t, *J* = 6.6 Hz, 2H), 2.18 – 2.13 (m, 2H), 1.89 (p, *J* = 6.3 Hz, 2H), 1.59 (s, 3H), 1.38 (s, 3H).

#### 4.2.3. General procedure for phthalidomides deprotection 5a-b

To a solution of **4a-b** (1 mmol, 1 equiv) in methanol (30 mL) was added hydrazine (0.1 ml, 3 mmol, 3 equiv). The reaction mixture was stirred at room temperature overnight. 10% acetic acid (3 ml) was added and extracted with ethylacetate (5 mL). 2 N sodium hydroxide (3 ml) was added and extracted with ethylacetate (3 x 5 mL). The combined organic extract was washed with brine, dried over anhydrous sodium sulfate, and the solvent was evaporated under *vacuo*. The residue was used in the next step without further purification.

#### 4.2.4. General procedure for amidation

To a solution of amino acid Fmoc-Orn(Mtt) or Fmoc-Lys(Boc) (0.3 mmol, 1 equiv) in acetonitrile (3 mL) was added ammonium chloride (53 mg, 0.6 mmol, 2 equiv), HBTU (379 mg, 0.45 mmol, 1.5 equiv), and N-methyl morpholine (105 mg, 0.6 mmol, 2 equiv). The reaction mixture was stirred at room temperature overnight. H_2_O was added (2 ml) and extracted with ethylacetate (3 x 7 mL). The combined organic extract was washed with brine, dried over anhydrous sodium sulfate, and the solvent was evaporated under *vacuo* and purified by flash chromatography on silica gel in 70-90% ethylacetate/hexanes to obtain **8a-b**.

#### 4.2.5. General procedure for peptide synthesis

The Mtt-protected ornithine/lysine-containing peptides **9a** and **9b** were synthesized using a Liberty Blue™ peptide synthesizer on Rink amide MBHA resin (138 mg with a resin loading of 0.1 mmol/g) using standard protocols with no final Fmoc deprotection. Each peptide was synthesized of 0.10 mmol and peptide couplings were performed by using 0.2 M of Fmoc-amino acid (Mtt-Orn or Mtt-Lys 2 equiv, Gly 6 equiv, Arg 12 equiv), activators of DIC reagent (4 ml of 0.5 M), and Oxyma (2 ml of 1 M) in DMF, 20% piperidine in DMF (13 ml) was used for deprotection. The resin was washed with DMF (3 x 3 mL) and DCM (2 x 2 mL). Peptides were not cleaved from the resin and used directly in the next step.

#### 4.2.6. General procedure for Mtt deprotection

To **8a, 9a-b** (0.1 mmol, 1 equiv) was added 1% trifluoroacetic acid in dichloromethane (1 ml). The reaction mixture was stirred at room temperature for 2 hr. The mixture was either evaporated under *vacuo* or washed with dichloromethane. The residue was neutralized using DIPEA (15 mg, 0.12 mmol, 1.2 equiv), which were used in the next step without further purification.

#### 4.2.7. Compound 8b Fmoc deprotection

To **8b** (0.3 mmol, 1 equiv) was added 20% trifluoroacetic acid in dichloromethane (3 ml). The reaction mixture was stirred at room temperature for 1 hr. The solvent was evaporated under *vacuo* and the residue was neutralized using DIPEA (45 mg, 0.36 mmol, 1.2 equiv) and the obtained residue was used in the next step without further purification.

#### 4.2.8. General procedure for synthesis of thiourea

To a solution of residues that obtained from Mtt and Fmoc deprotection in dichloromethane (3 mL) at 0°C was added Fmoc-isothiocyanate (42 mg, 0.15 mmol, 1.5 equiv). The reaction mixture was stirred at 0°C for 30-60 min. The mixture was evaporated under *vacuo* or washed. The evaporated residue was crystalized using dichloromethane:hexanes 1:1 (2 ml) to produce **6a-d**.

#### 4.2.9. General procedure for synthesis of guanidine moiety

To a solution of amine **5a-b** (0.05 mmol, 1 equiv) in dichloromethane (2 mL) was added thiourea **6a-d** (0.75 mmol, 1.5 equiv), EDC (20 mg, 0.1 mmol, 2 equiv), and DIPEA (13 mg, 0.1 mmol, 2 equiv). The reaction mixture was stirred at room temperature overnight. The mixture were washed with dichloromethane (3 x 2 mL) to produce **7a-h**.

#### 4.2.10. General procedure for final compounds synthesis

The mixture **7a-h** were treated with 20% piperidine in DMF with 0.1 M HOBt (3 x 2 mL) for 10 min each. Then it was flushed with dichloromethane (3 x 2 mL) and used for next step. Then it was dissolved in dichloromethane (2 mL) followed by adding a cleavage cocktail (TFA:TIPS:DODT:H_2_O) (9.4 mL:0.1 mL:0.25 mL:0.25 mL). The reaction mixture was stirred at room temperature for 5 hr. The mixture was evaporated by passing N2 gas, washed with dry ether. The residue was used for semi-preparatory HPLC separation using MeOH/H_2_O 10-40% to obtain final compounds **1a-h**.

##### 4.2.10.1. AH240 1a (Lys-C2-AD)

C_19_H_32_N_10_O_4_S, MALDI, calcd for [M + H]^+^ 497.2407; found 497.3052.

##### 4.2.10.2. AH244 1b (Lys-C3-AD)

C_20_H_34_N_10_O_4_S, MALDI, calcd for [M + H]^+^ 511.2563; found 511.2719.

##### 4.2.10.3. AH238 1c (Arg-C2-AD)

C_18_H_30_N_10_O_4_S, MALDI, calcd for [M + H]^+^ 483.2172; found 483.2628.

##### 4.2.10.4. AH246 1d (Arg-C3-AD)

C_19_H_32_N_10_O_4_S,_MALDI, calcd for [M + H]^+^ 497.2407; found 497.2405.

##### 4.2.10.5. AH218 1e (RGK-C2-Ad)

C_26_H_45_N_15_O_6_S,_MALDI, calcd for [M + H]^+^ 710.3554; found 710.6207.

##### 4.2.10.6. AH221 1f (RGK-C3-Ad)

C_28_H_49_N_15_O_6_S,_MALDI, calcd for [M + H]^+^ 724.3711; found 724.5140.

##### 4.2.10.7. AH233 1g (RGR-C2-Ad)

C_26_H_45_N_15_O_6_S, MALDI, calcd for [M + H]^+^ 696.3398; found 696.4561.

##### 4.2.10.8. AH237 1h (RGR-C3-Ad)

C_27_H_47_N_15_O_6_S, MALDI, calcd for [M + H]^+^ 710.3554; found 710.4666.

### 4.3. The Enzyme Inhibition Assay and Selectivity Studies

A fluorescence-based SAHH-coupled assay was employed to calculate their IC_50_ values and to study the effect of the synthesized compound on methyltransferase activity of PRMT1, G9a, SETD7, NTMT1, and *Tb*PRMT7.^21,22^ For PRMT1, the assay was carried out in a final well volume of 40 μL: 25 mM HEPES buffer (pH = 8), 25 μM EDTA, 25 mM NaCl, 0.01% Triton X-100, 50 μM TCEP, 5 μM SAHH, 0.1 μM PRMT1, 5 μM SAM, and 15 μM ThioGlo1. The inhibitor was added at nine compound concentrations: 0.005, 0.015, 0.046, 0.14, 0.41, 1.23, 3.7, 11, and 33 μM. After 10 min incubation with the inhibitor, reactions were initiated by the addition of 4 μM H4-21 peptide. For G9a, the assay was performed in a final well volume of 40 μL: 25 mM potassium phosphate buffer (pH = 7.6), 1 mM EDTA, 2 mM MgCl_2_, 0.01% Triton X-100, 5 μM SAHH, 0.1 μM G9a, 10 μM SAM, and 15 μM ThioGlo1. The inhibitor was added at a single compound concentration of 100 μM. After 10 min incubation with the inhibitor, reactions were initiated by the addition of 4 μM H3-21 peptide. For SETD7, the assay was performed in a final well volume of 40 μL: 25 mM potassium phosphate buffer (pH = 7.6), 0.01% Triton X-100, 5 μM SAHH, 1 μM SETD7, 2 μM AdoMet, and 15 μM ThioGlo1. The inhibitors were added at a single compound concentration of 100 μM. After 10 min incubation with inhibitors, reactions were initiated by the addition of 90 μM H3-21 peptide. For NTMT1, the assay was exerted in a final well volume of 40 μL: 25 mM Tris (pH = 7.5), 50 mM KCl, 0.01% Triton X-100, 5 μM SAHH, 0.2 μM NTMT1, 3 μM SAM, and 15 μM ThioGlo1. The inhibitor was added at a single compound concentration of 100 μM. After 10 min incubation with the inhibitor, reactions were initiated by the addition of 3 μM SPKRIA peptide. For *Tb*PRMT7, the assay was performed in a final well volume of 40 μL: 25 mM Tris (pH = 7.5), 50 mM KCl, 0.01% Triton X-100, 5 μM SAHH, 0.2 μM *Tb*PRMT7, 3 μM SAM, and 15 μM ThioGlo1. After 10 min incubation with inhibitors, reactions were initiated by the addition of 60 μM H4-21 peptide. Fluorescence was monitored on a BMG CLARIOstar microplate reader with excitation 400 nm and emission 465 nm. Data were processed by using GraphPad Prism software 7.0.

The HotSpot methyltransferases profiling (radioisotope filter binding platform) was used to evaluate the selectivity of compound **AH237** against 41 methyltransferases (Reaction Biology Corp.). The compound was tested in single dose (10 μM) with duplicate. Control compounds, SAH (S-(5’-Adenosyl)-L-homocysteine), Chaetocin, LLY 507, or Ryuvidine were tested in 10-dose IC_50_ mode with 3-fold serial dilution starting at 100 or 200 μM. All the reactions were carried out at 1 μM SAM.

### 4.4. The Enzyme Inhibition Study

For the enzyme kinetic study, the fluorescence-based SAHH-coupled assay was used. Varying concentrations of SAM (from 0.5 to 16 Km) with 60 μM fixed concentration of H4-21 or varying concentration of H4-21 (from 0.5 to 12 Km) with 3 μM fixed concentration of SAM were included in reactions with a series concentration of compounds. All the IC_50_ values were determined in triplicate. Fluorescence was monitored on a BMG CLARIOstar microplate reader with excitation 380 nm and emission 505 nm. Data were processed by using GraphPad Prism software 7.0.

### 4.5. MS Methylation Inhibition Analysis

MALDI-MS methylation inhibition assay was performed and analyzed via a Sciex 4800 MALDI TOF/TOF MS. 0.2 μM TbPRMT7, 20 mM Tris (pH = 7.5), 50 mM KCl, 5 μM AdoMet and various concentrations of compounds at 37 °C for 10 min before the addition of 2.5 μM H4-21 peptide to initiate the reaction. After incubation overnight, the samples were quenched in a 1:1 ratio with a quenching solution (20 mM NH4H2PO4, 0.4% (v/v) TFA in 1:1 acetonitrile/water). Samples were analyzed by MALDI-MS with 2,5-dihydroxybenzoic acid matrix solution. Duplicate was performed for all experiments. Data were processed in Data Explorer.

### 4.6. Molecular Modeling

Glide of Schrödinger Maestro (Version 10.1) was used to predict the binding of **AH237** against the active site of PRMT 1, 4, 5, and 7. The chemical structure of **AH237** was drawn with Ligprep module and ionization state was generated at pH 7.0 ± 2.0 using Epik module. PRMT1 (1OR8), PRMT4 (5LGP), PRMT5 (4GQB), and PRMT7 (4M38) were obtained from the Protein Data Bank. Structures were prepared and refined with the protein preparation module and the energy was minimized using OPLS_2005 force field. Grids were generated with Glide by adopting the default parameters. A cubic box of specific dimensions centered around the active site residues was generated for the receptors. The bounding box was set to 15 Å × 15 Å × 15 Å. Flexible ligand docking was performed. Glide extra precision docking was performed by keeping all docking parameters as default. Ligand poses were generated for **AH237** by using Monte Carlo random search algorithm, and its binding affinities to PRMTs were predicted with Glide docking score. Post-docking minimizations were taken under OPLS_2005 force field, and 30 poses per ligand were finally saved.

## Acknowledgments

The authors acknowledge the support from NIH grants R01GM117275 (RH) and P30 CA023168 (Purdue University Center for Cancer Research). We also thank supports from the Department of Medicinal Chemistry and Molecular Pharmacology (RH) at Purdue University.

## AUTHOR INFORMATION

Corresponding Author. * Phone: (765) 494 3426.

## Author contributions

The manuscript was written through contributions of all authors. All authors have given approval to the final version of the manuscript.

## Conflicts of interest

The authors declare no competing financial interest.

## Appendix A. Supporting information

Supporting data to this article can be found online

## Supporting Information

**Table S1.** The single dose of HotSpot methyltransferases inhibition for a single dose (10 μM) of **AH237** on various methyltransferases. These data are associated with Figure 3 in the manuscript.

| **Methyltransferase** | **Enzyme % Activity\*** | **Methyltransferase** | **Enzyme % Activity\*** |
| --- | --- | --- | --- |
| **ASH1L** | 100.10 | **NSD2 (E1099K)** | 107.95 |
| **DNMT1** | 94.27 | **NSD2 (T1150A)** | 108.71 |
| **DNMT3a** | 86.34 | **NSD3** | 102.65 |
| **DNMT3b** | 88.39 | **PRDM9** | 110.83 |
| **DNMT3b/3L** | 103.47 | **PRMT1** | 28.96 |
| **DOT1L** | 74.89 | **PRMT3** | 100.72 |
| **EZH1 Complex** | 92.13 | **PRMT4** | 0.28 |
| **EZH2 Complex** | 97.83 | **PRMT5/MEP50 Complex** | 0.20 |
| **EZH2(Y641F)**<br>**Complex** | 95.23 | **PRMT5 (C449S)/MEP50** | 0.70 |
| **G9a** | 88.07 | **PRMT6** | 41.59 |
| **GLP** | 103.64 | **PRMT7** | 25.08 |
| **METTL21A** | 97.44 | **PRMT8** | 59.64 |
| **MLL1 Complex** | 111.76 | **SET1b Complex** | 96.84 |
| **MLL2 Complex** | 110.15 | **SET7/9** | 110.28 |
| **MLL3 Complex** | 102.61 | **SET8** | 99.68 |
| **MLL4 Complex** | 97.85 | **SETD2** | 91.70 |
| **NTMT1** | 102.22 | **SMYD2** | 50.67 |
| **NTMT2** | 91.14 | **SMYD3** | 50.28 |
| **NSD1** | 85.71 | **SUV39H1** | 100.32 |
| **NSD2** | 101.46 | **SUV39H2** | 100.12 |
|  |  | **SUV420H1TV2** | 96.69 |

\*relative to DMSO control

### ^1^H NMR and ^13^C spectra

***S*-(((3a*S*,4*S*,6*R*,6a*R*)-6-(6-Amino-9*H*-purin-9-yl)-2,2-dimethyltetrahydrofuro[3,4-*d*][1,3]dioxol-4-yl)methyl) ethanethioate (2)**

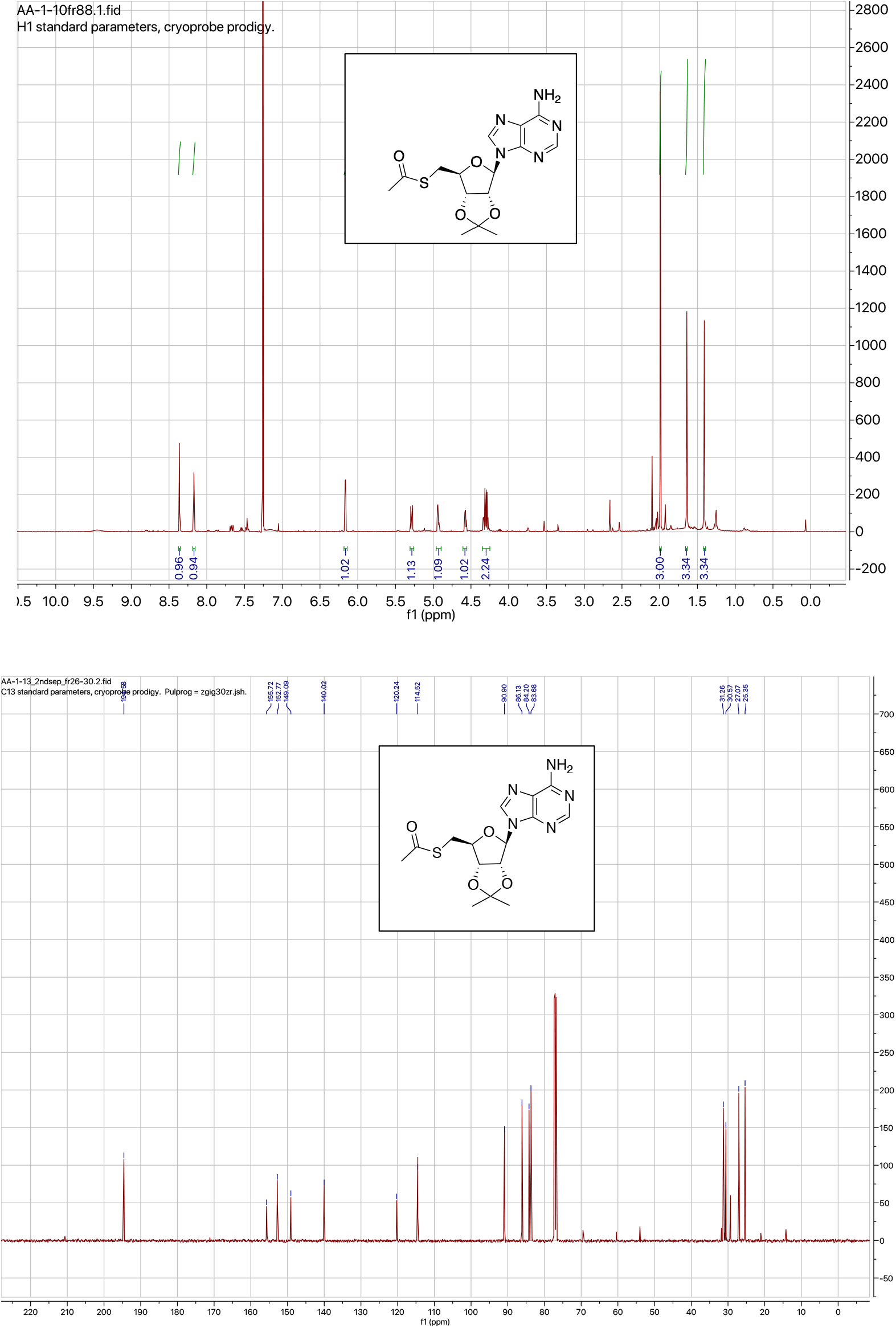

**2-(2-((((3a*S*,4*S*,6*R*,6a*R*)-6-(6-Amino-9*H*-purin-9-yl)-2,2-dimethyltetrahydrofuro[3,4-*d*] [1,3]dioxol-4-yl)methyl)thio)ethyl)isoindoline-1.3-dione (4a)**

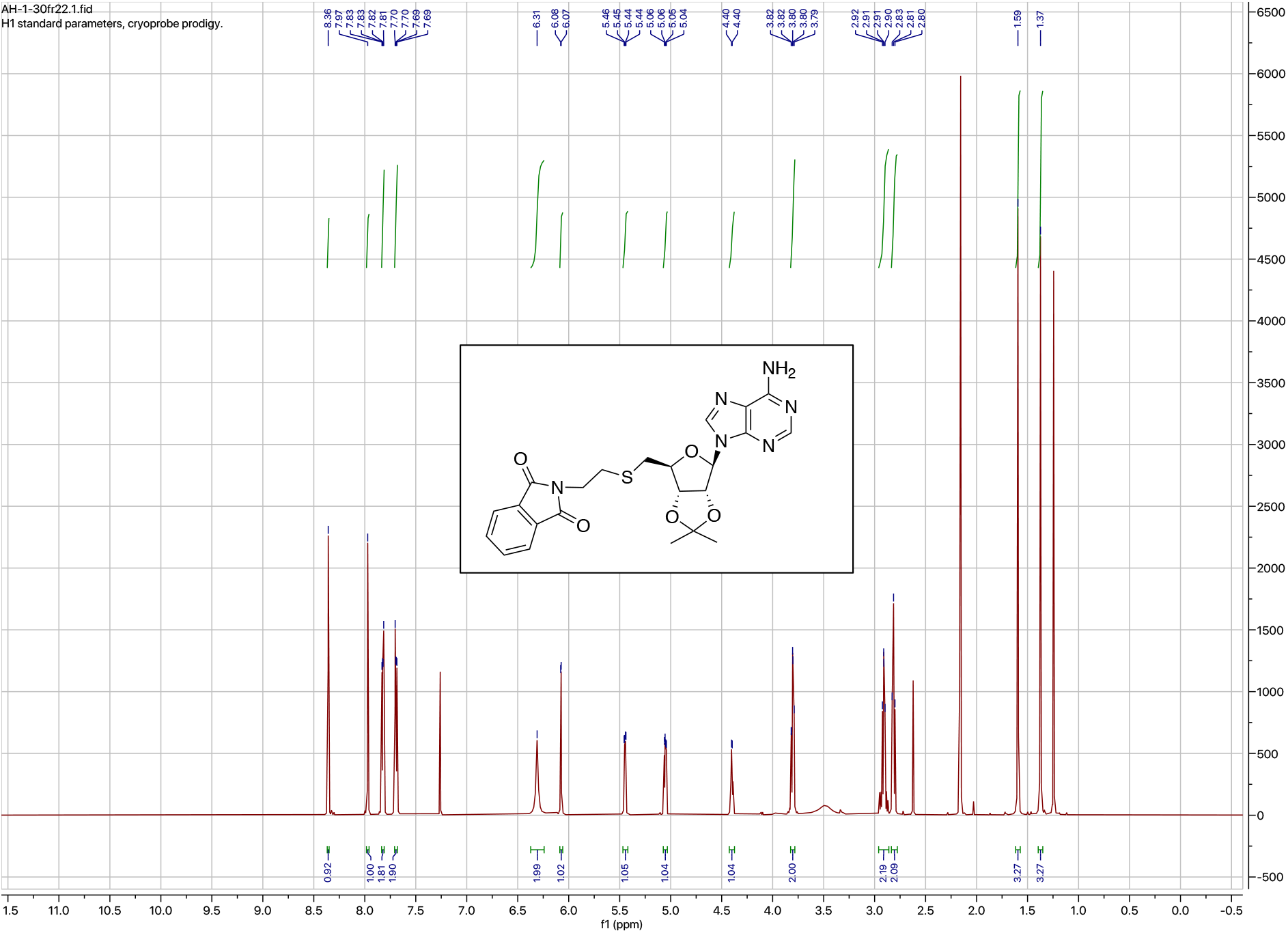

**2-(3-((((3a*S*,4*S*,6*R*,6a*R*)-6-(6-Amino-9*H*-purin-9-yl)-2,2-dimethyltetrahydrofuro[3,4-*d*][1,3]dioxol-4-yl)methyl)thio)propyl)isoindoline-1,3-dione (4b)**

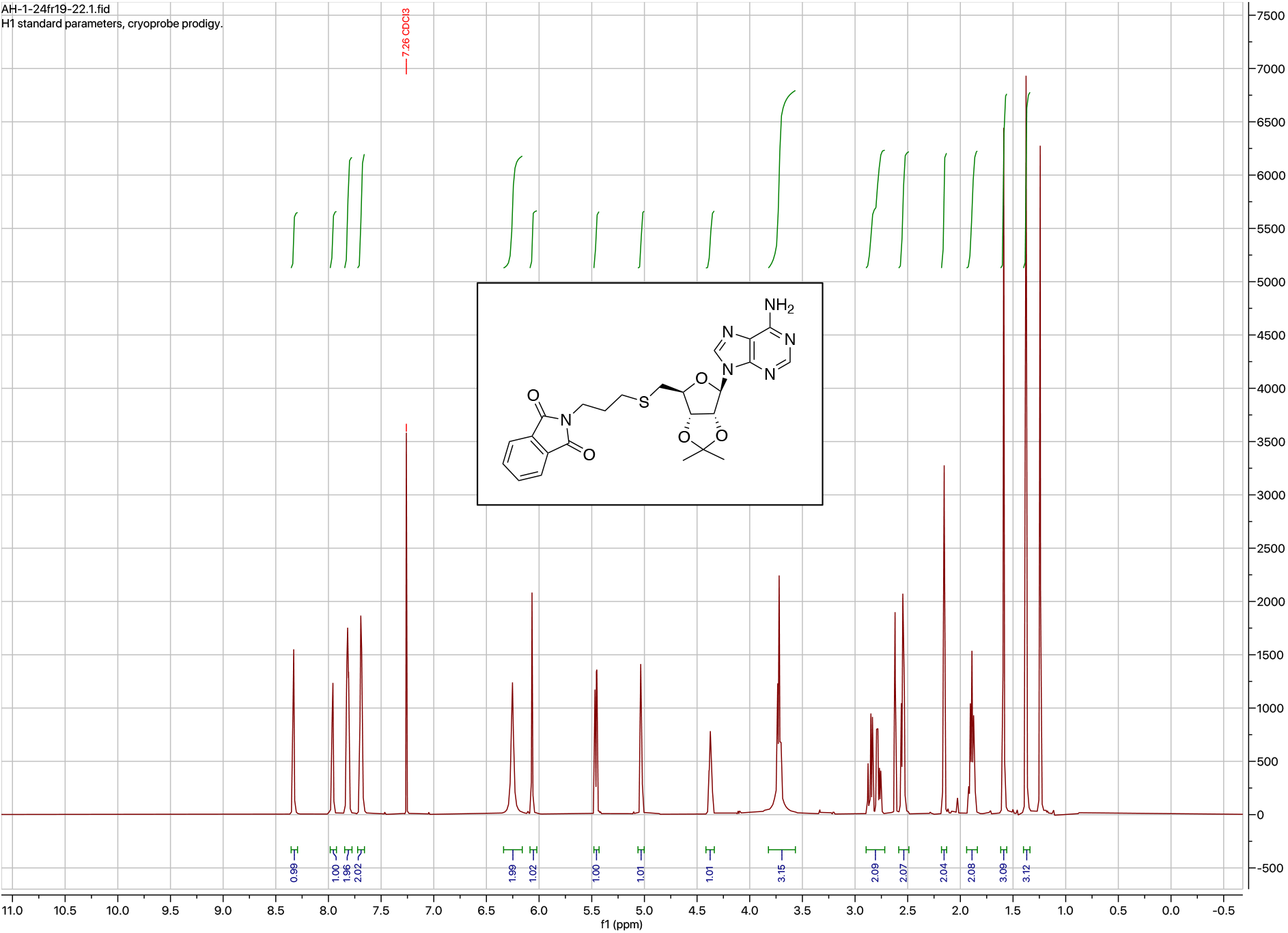

### MALDI spectra

**AH240 (1b) (Lys-C2-Ad).**

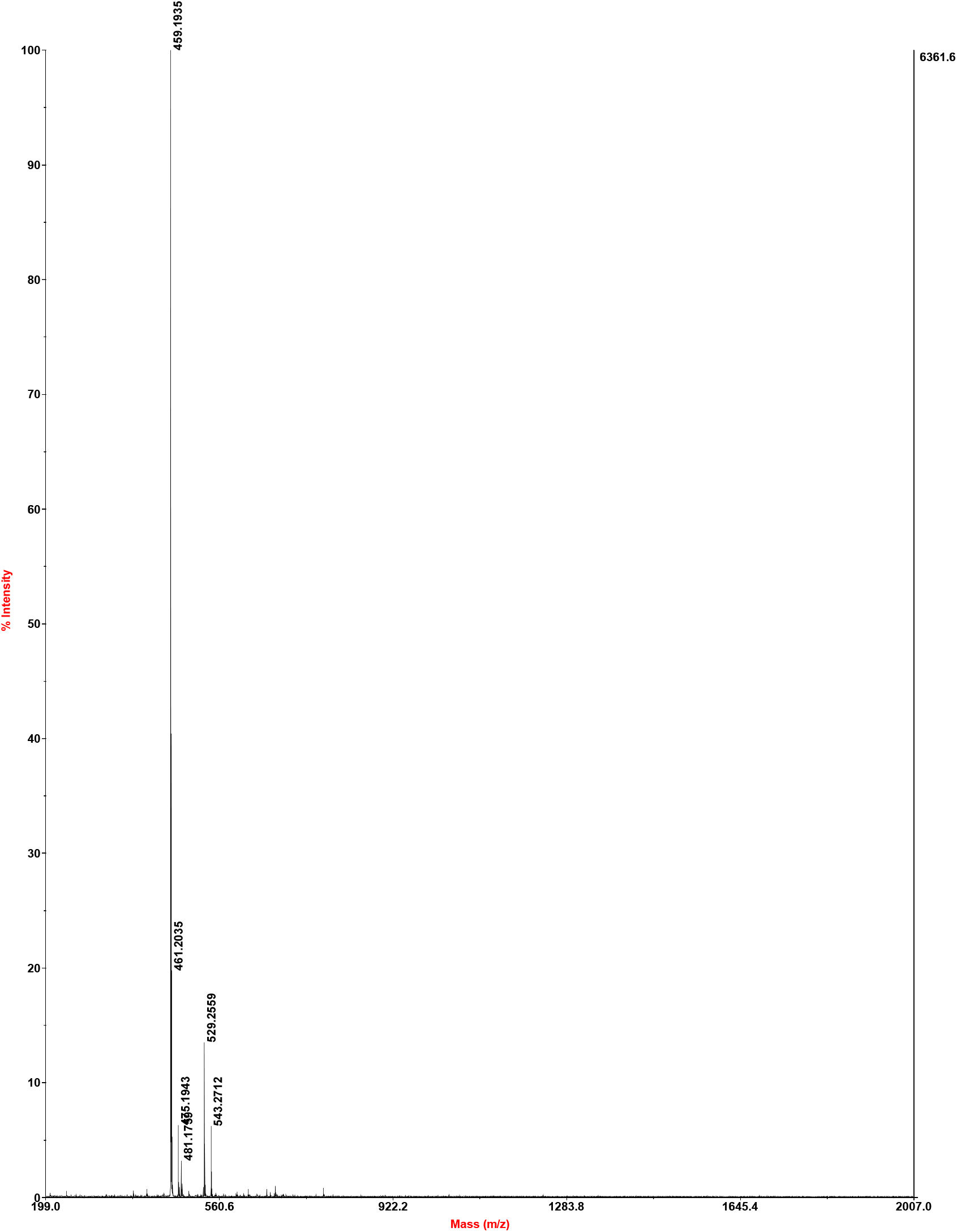

**AH244 (1f) (Lys-C3-Ad).**

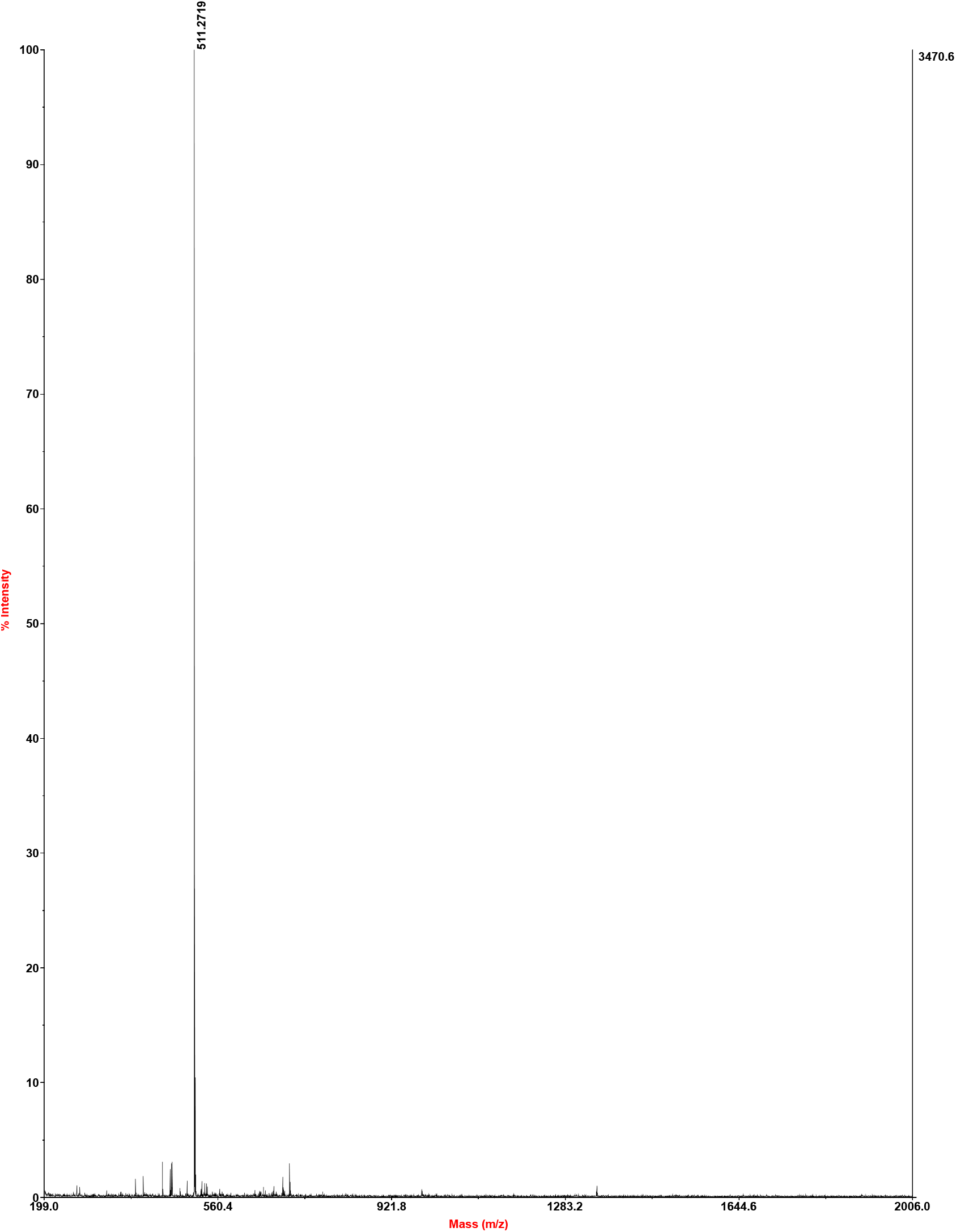

**AH238 (1a) (Arg-C2-Ad).**

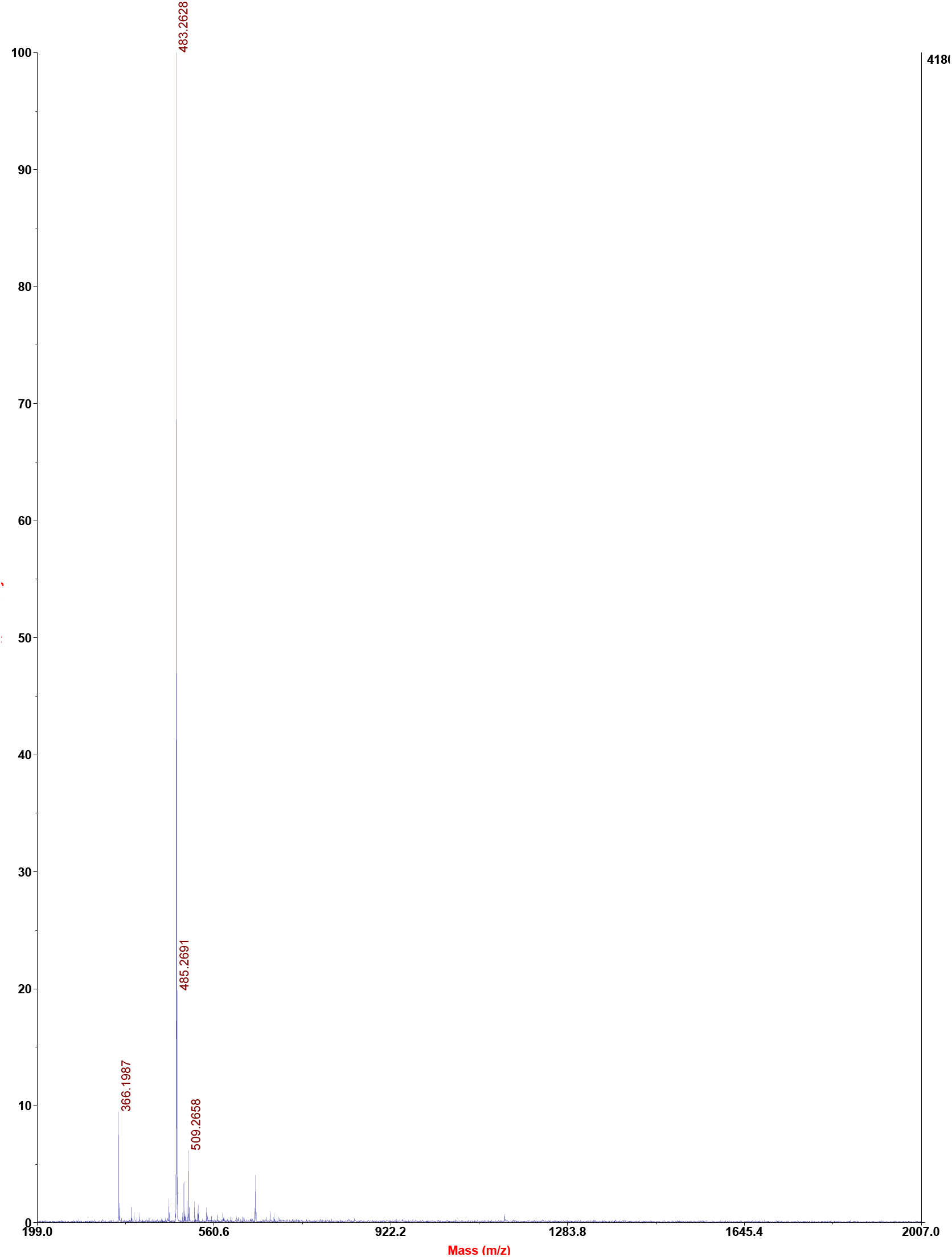

**AH246 (1e) (Arg-C3-Ad).**

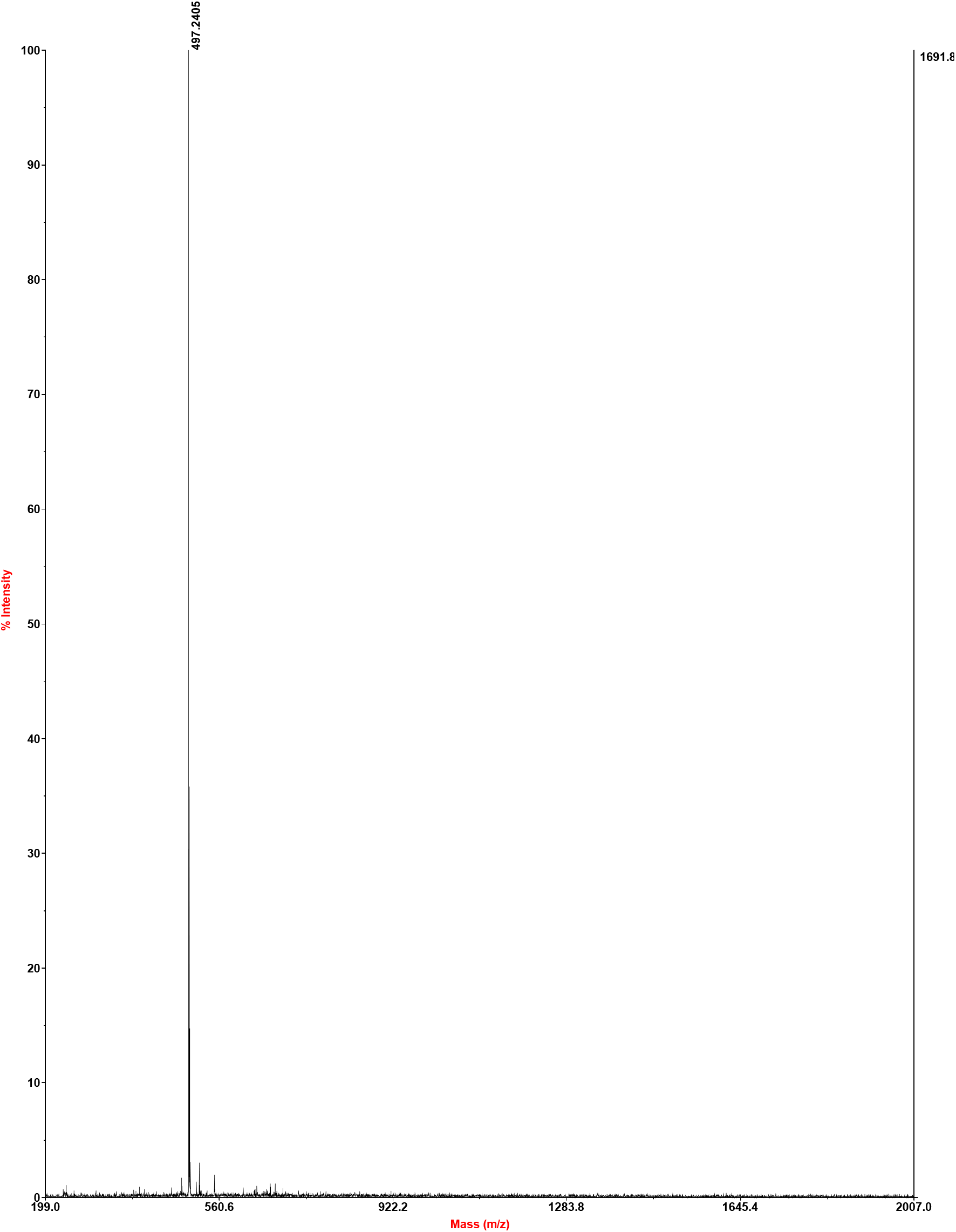

**AH218 (1d) (RGK-C2-Ad).**

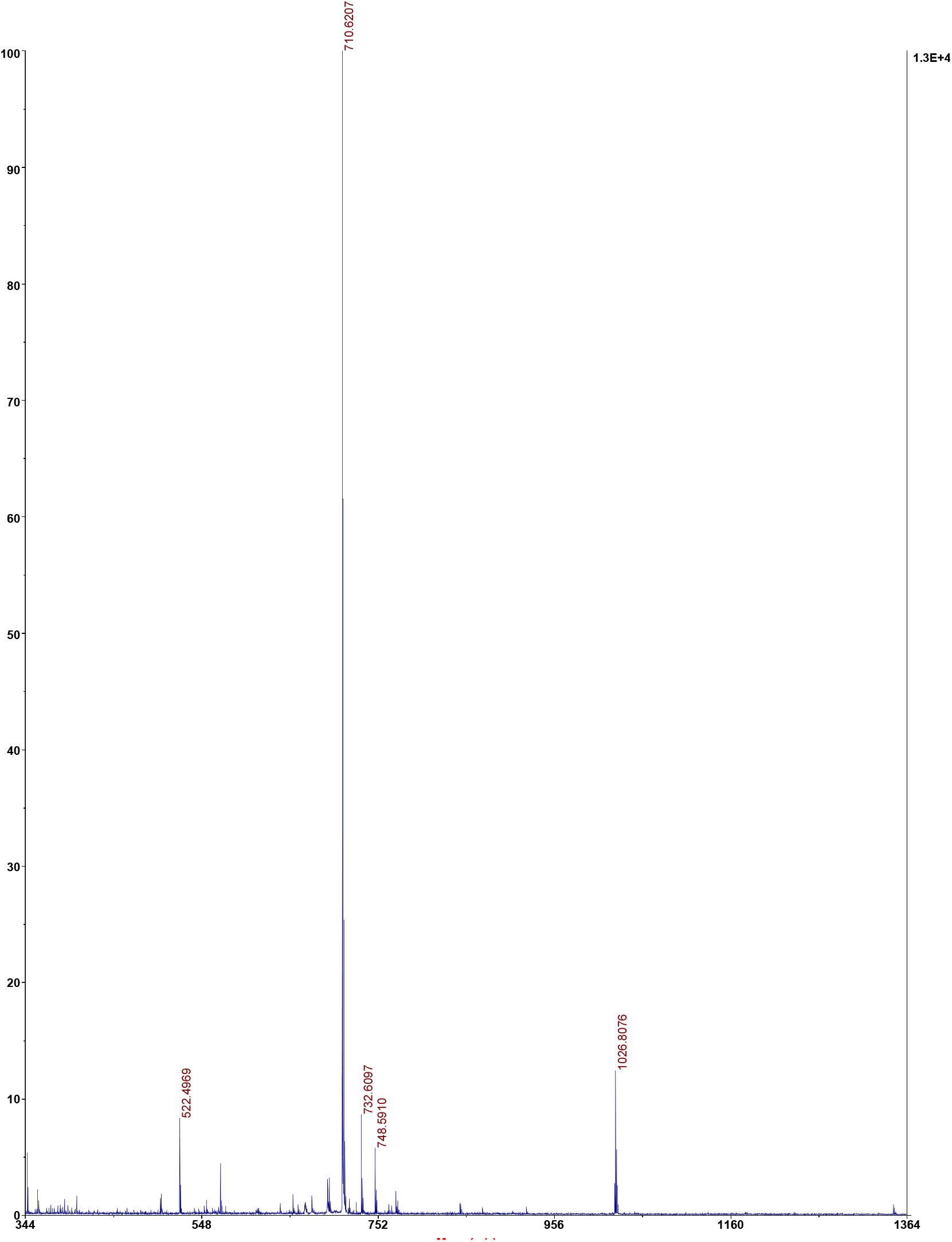

**AH221 (1h) (RGK-C3-Ad).**

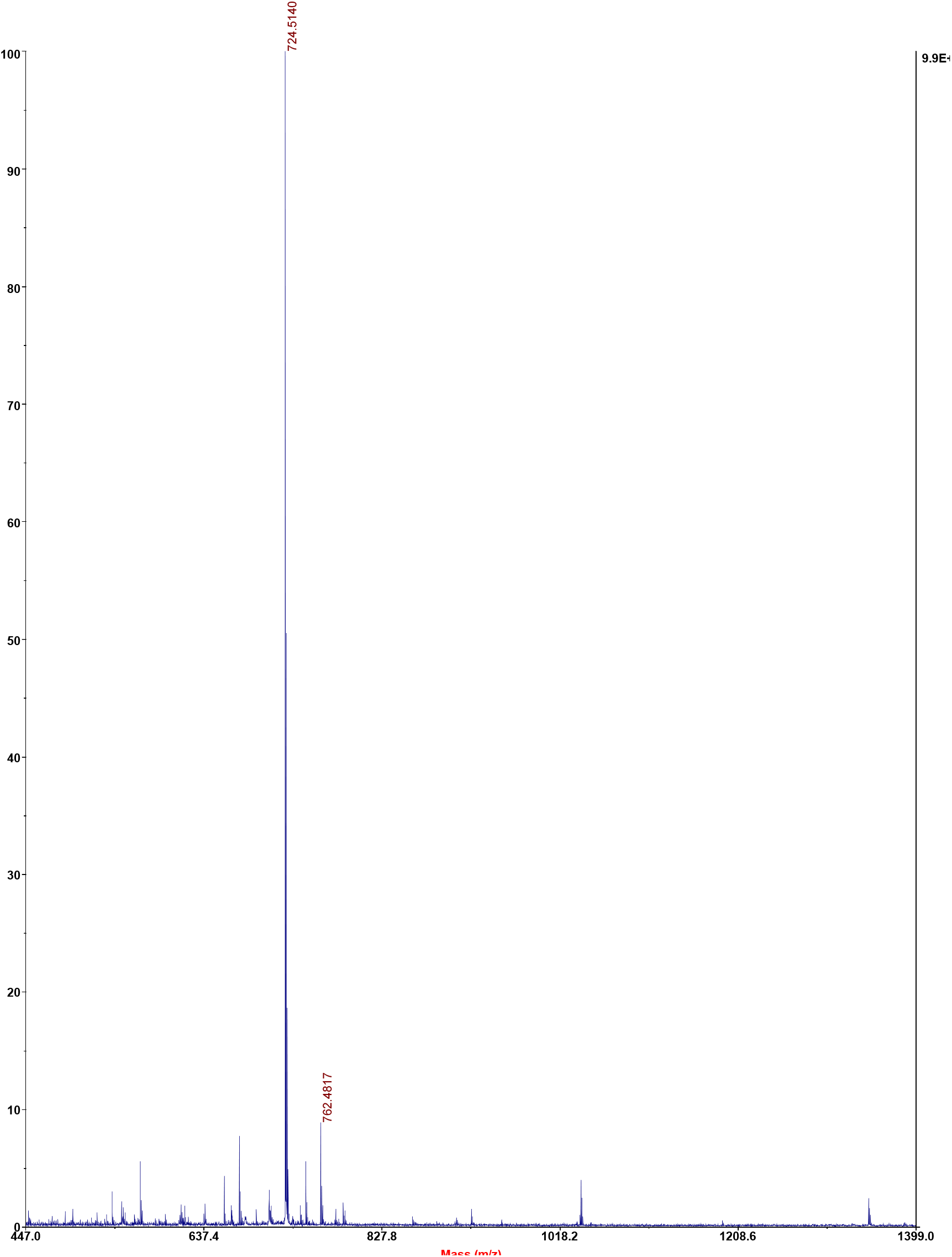

**AH233 (1c) (RGR-C2-Ad).**

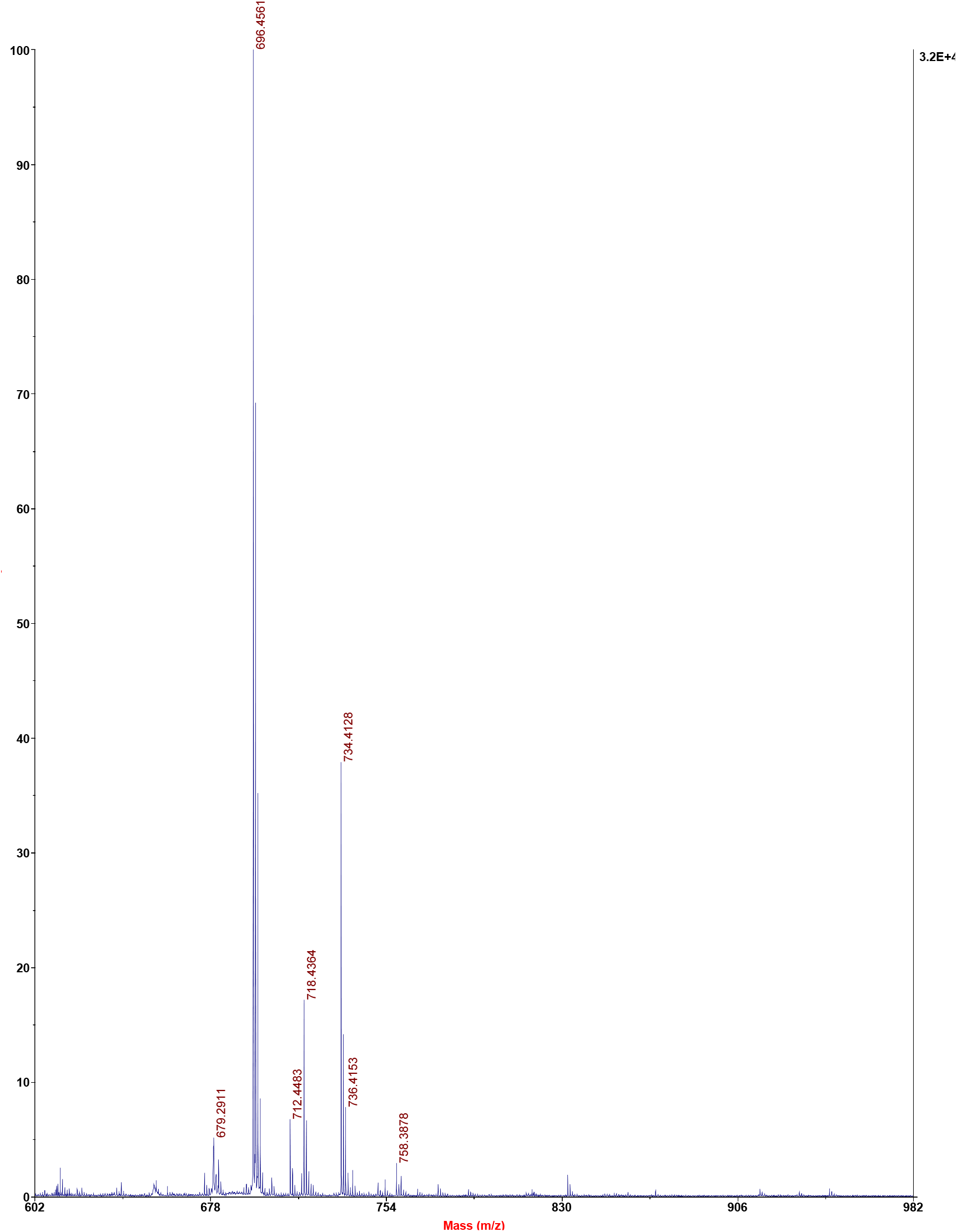

**AH237 (1g) (RGR-C3-Ad).**

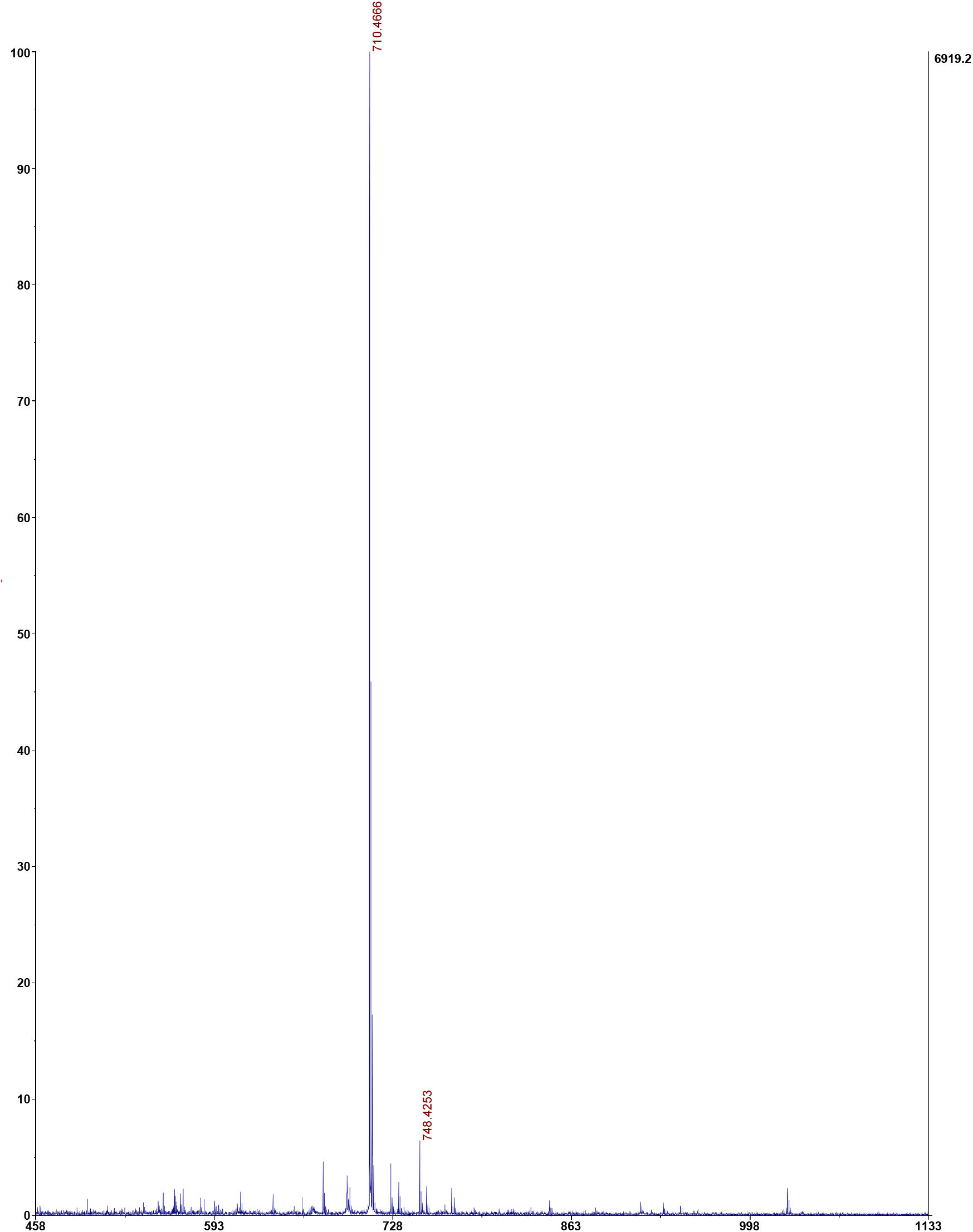

## Notes

### Competing Interest Statement

The authors have declared no competing interest.

